# Originality in creative ideation evolves along nonlinear axes in the human brain

**DOI:** 10.1101/2025.06.03.657648

**Authors:** Rikki Rabinovich, Rhiannon L. Cowan, Mark R. Libowitz, Tyler S. Davis, Shervin Rahimpour, Roger E. Beaty, Eleonora Bartoli, Sameer A. Sheth, Elliot H. Smith, Ben Shofty

## Abstract

The ability to think creatively is fundamental to human cognition, shaping how we interact with the world and navigate complex problems. Creativity depends on the activity of multiple large-scale brain networks, but the neural representations that underlie creative thought and distinguish it from other cognitive states remains unclear. Here, we use multiscale intracranial electrophysiology—from single-unit recordings to local field potentials across distributed brain networks—to discover how the human brain generates creative ideas. Our results demonstrate that cortical networks in the human brain display activity patterns that differentiate creative thinking from non-creative cognitive processes such as arithmetic calculations. We found that the underlying geometry of neural activity in the default mode network is shaped by instantaneous cognitive state. Further, in the dorsal attention network, we uncovered nonlinear, high-dimensional representations of moment-by-moment fluctuations in creative performance. These representations define a neural creativity axis that is shared between network-level and single-neuron computations. Our findings reveal widespread neural representations of cognitive state and suggest distinct roles of specific cortical networks in controlling creativity, with the default mode network enabling creative states, while higher-dimensional dorsal attention network activity, associated with attentional selection or other context-dependent constraints, reflects the production of more original responses.

## Introduction

The capacity for creative thought is central to human cognition, guiding our approach to complex challenges. Among forms of internal mentation, creativity is unique in that it involves a fluid, multimodal interaction between introspection and the external world (e.g., generating novel solutions to existing problems). While creativity encompasses a multifaceted set of processes, here, we use the term to specifically refer to divergent thinking. Divergent thinking is defined operationally as the internal generation of multiple, non-obvious, and conceptually distant ideas^1,2^. In particular, the present study focuses on idea originality (or conceptual novelty) as a measurable facet of creative cognition.

As a vital component of human experience and essential to societal progress, creativity has received considerable attention from the scientific community; yet, despite numerous attempts to characterize this cognitive process, the neural underpinnings of creative thinking remain elusive. So far, prior research has led to a consensus that no single brain region is responsible for divergent thinking; rather, activity patterns of large-scale cortical networks linking widespread brain areas likely underlie creative ideation and related cognitive processes^3–7^.

One such network, known as the default mode network (DMN), is thought to play a role in self-referential or internally-directed cognition. Originally conceptualized as the default mode of the brain, this network is active during periods of rest and is often suppressed by task engagement and external attention^8^. Since then, it has been implicated in cognitive processes ranging from episodic memory to dreaming to creative thinking^9,10^. Specifically, existing literature links DMN activity to the generation of novel ideas, though the contribution of DMN to creativity likely depends on interaction with cognitive control networks, which may regulate and filter creative output to ensure generated ideas are both original and useful^11–15^. However, most prior studies focus on individual differences in creativity or static measures of network engagement, without resolving the neural dynamics that define creative states within individuals^9,16–18^. Further, while one causal study found that disrupting the DMN did not affect the quality or originality of responses—only the number generated^11^—another found that stimulating the DMN did impact originality^19^. This discrepancy underscores the complexity of creative ideation and reveals the lack of a unifying model for divergent thinking. Thus, a fundamental question remains: what distinguishes creative thought in the brain and how do neural dynamics support it?

The present study combines invasive recordings in the human brain with machine learning and dimensionality reduction approaches to examine neural representations of creativity at multiple scales, ranging from single-neuron activity to large-scale brain networks, with the goal of uncovering the representational geometries underlying creative ideation. We combine this analysis of representational geometry with the temporal resolution of intracranial electrophysiology to read out and predict ongoing moment-by-moment fluctuations in cognitive states, something that would not be possible with non-invasive neuroimaging. Thus, we aimed to directly address questions about state geometry, dimensionality, and cross-scale consistency, using out-of-sample prediction and cross-validation to move beyond traditional correlational frameworks.

We hypothesized that DMN activity patterns would contain distinct representations of the creative state as well as encode the quality of creative output. We predicted that other networks may become engaged at various points in the task but would only weakly encode creativity. Finally, we hypothesized that creativity representations could be identified at the single neuron level. While our data supported some of our hypotheses, our results painted a much more complex picture of the network-level encoding of creative thought.

## Results

To probe the role of the DMN and other networks in divergent thinking, we leveraged intracranial electrodes implanted in the brains of 17 neurosurgical patients undergoing monitoring for drug-resistant epilepsy; from these electrodes, we directly recorded neural signals in the human brain while subjects either engaged in creative thinking or other forms of thought (computing simple arithmetic or describing object characteristics).

To distill creative ideation into a simple process that can be analyzed and manipulated, we applied a version of the Alternate Uses Task (AUT). This paradigm is a widely used method for probing divergent thinking and requires participants to invent creative uses for everyday objects^2,3,16,17^. A Basic Arithmetic Task (BAT), in which participants solved simple addition or subtraction problems, served as our main contrast (Fig. 1a). We chose BAT as our comparison task based on precedents in prior research where arithmetic calculations were used as control conditions for verbal tasks and because of shared semantic control processes across both verbal and numerical domains^20,21^. Both tasks had identical structure (silent stimulus window followed by spoken verbal response) to ensure that the only distinction between the tasks was the cognitive component. However, to account for possible unintended differences between AUT and BAT unrelated to creativity, a subset of patients (n=9) also completed an Object Characterization Task (OCT), which asked participants to report the most defining characteristic of the presented object. The OCT, like AUT, is semantic, open-ended, and internally directed due to the engagement of autobiographical memory, but does not prompt the participant to think divergently.

**Figure 1.**
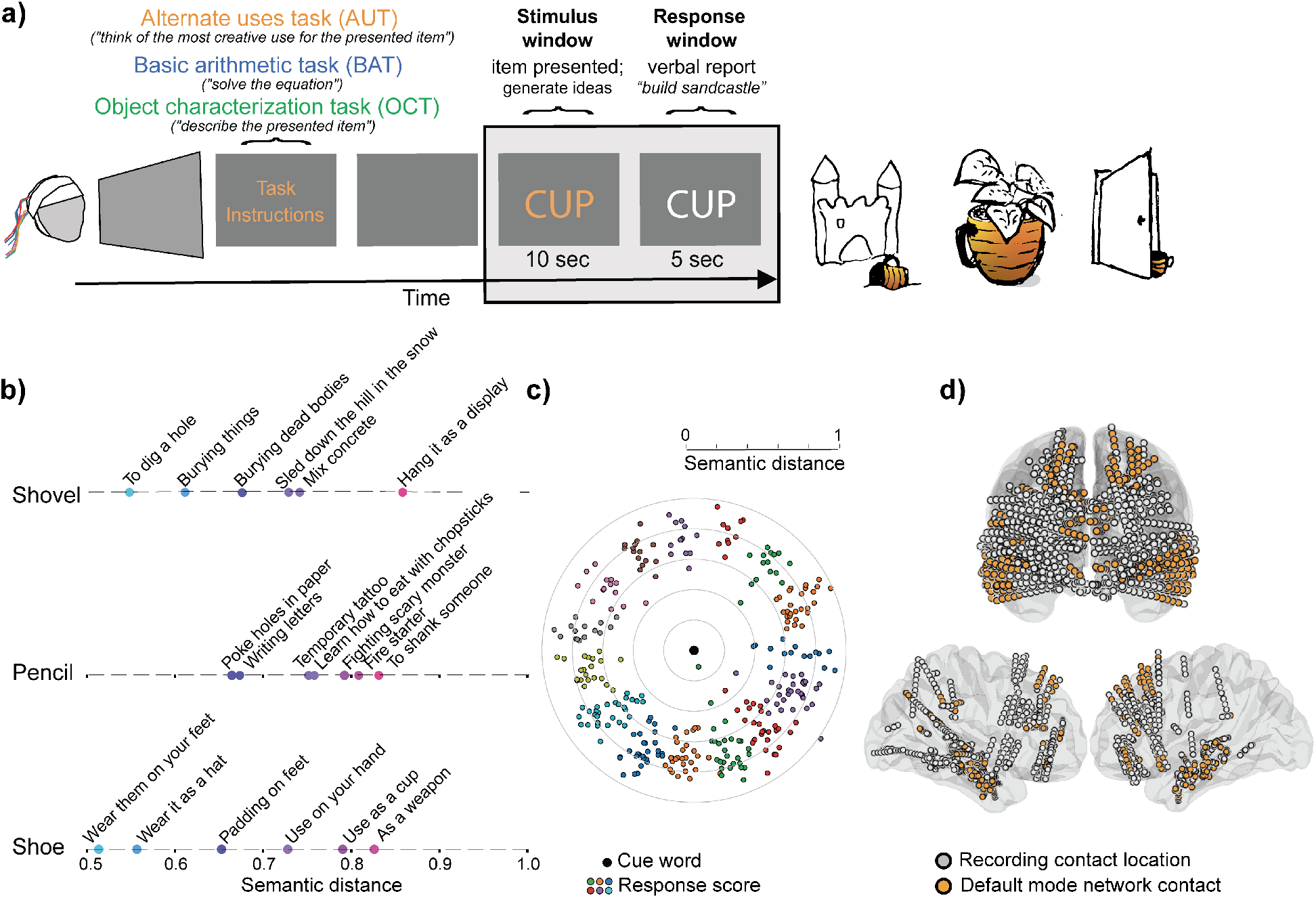
Experimental design. **A)** Left: task design. Trials occur in blocks of three, with instructions appearing on the screen at the start of each block. For each trial, the prompt (in a colored font) is present for a 10 second stimulus window, after which the font turns white, indicating the beginning of the 5 second response window. Right: stylized example responses. For the prompt “cup”, possible high-originality uses include “build a sandcastle” (semantic distance score of 0.909), “turn it into a planter” (0.930), or “use as a doorstop” (0.967). (**B**) Patients’ responses for 3 example words. **C)** All patients’ responses across all trials (n=15 patients; 316 trials with valid responses). Each dot represents a single trial’s response; distance from center is the semantic distance score for that response. Each cluster of same-color dots corresponds to 1 patient. **D)** Signals are recorded from implanted electrodes during task execution. The model brain shows locations of the full set of contacts for all patients (n=17). Linearly aligned contacts resided on the same electrode lead; contacts on each depth electrode that were classified as belonging to the DMN are colored orange. Note that electrode contact localization required warping from patient space to MNI space due to anatomical variations between subjects, while Figure 1d overlays all electrodes on an average transparent model brain. Thus, the goal of Figure 1d is to illustrate approximate locations of recording sites, not to show precise anatomical placement.

In our version of the AUT (Fig. 1a), a cue word (e.g., “cup”) appeared on a screen: during the 10-second stimulus window, subjects silently thought of possible uses for the item. Then, a change of font color prompted subjects to choose a single most creative use for the object and verbalize their response. Responses were automatically scored for originality using natural language processing tools (semantic distance, Fig. 1b). For the prompt “cup”, the response “drink tea” scores low on originality (semantic distance=0.504), while “build a sandcastle” scores high (0.943), as it is semantically far from the meaning of the cue word. Trials were presented in blocks of three, such that each block consisted of three sequential trials of one type, before moving on to another trial type.

Importantly, we do not assume that participants’ intuitive judgments of creativity are identical to semantic distance; indeed, subjective creativity judgments are known to be variable across individuals and evaluators. Instead, semantic distance provides a standardized operationalization of originality, reflecting the novelty of responses, and has been shown to correlate with human ratings of creativity while being more reliable^22^. Thus, semantic distance is an accepted measure of quantifying creativity^15,19,22–25^.

Patients exhibited satisfactory task performance; each subject’s set of responses spanned a range of semantic distance scores, indicating trial-to-trial variability in creativity level (Fig. 1b; mean originality=0.716; standard deviation=0.123); meanwhile, the distribution of scores was similar across patients, with no outlier subjects. Among the datasets presented here, we never observed nonsensical responses, in which the answer was completely unrelated to the prompt word. During task execution, we recorded LFP and single-unit responses from intracranial depth electrodes implanted throughout various brain regions and cortical networks (Fig. 1c). Networks sampled were default mode network (DMN), frontoparietal network (FPN), somatomotor network (SMN), limbic network, visual network, dorsal attention network (DAN), and ventral attention networks (VAN).

### Network dynamics reflect cognitive state

First, we examined the temporal dynamics of DMN engagement in the creativity task as well as the control tasks (Fig. 2, Fig. 3a,b) to ensure reproducibility both of our prior observation of DMN recruitment in creativity^19^ and of previously established canonical disengagement of DMN in attention-demanding tasks like our arithmetic task^26,27^. We found that creativity trials displayed qualitatively increased high-frequency broadband activity (70-140Hz, also referred to as high gamma) and decreased low-frequency (theta; 4-8Hz) LFP power, relative to baseline (computed as the average of all intervals 2 – 1 seconds before the start of each block consisting of three trials; see Supp. Fig. 1a and Methods) and to control tasks. Specifically, during AUT, high gamma was elevated, exhibiting the greatest increase during the response window (aligning LFP activity to speech onset yielded qualitatively similar results as aligning to defined trial windows; Supp. Fig. 1b). Cluster-based permutation tests revealed a significant increase relative to baseline from 1 – 5 seconds into the response window, with a mean increase of 27.41% (*p*=0.0050, cluster-corrected across the 5 second response window). The fact that DMN recruitment (as measured by high gamma power in AUT) was strongest during the response period is consistent with our prior results in a different version of AUT^19^; however, interpretation of response-period effects merits caution, as these effects may reflect a mixture of ongoing idea generation, delayed internal decision timing, speech preparation, and verbal report. Low-frequency power, by contrast, decreased in DMN during both the stimulus and response periods of the creativity task.

**Figure 2.**
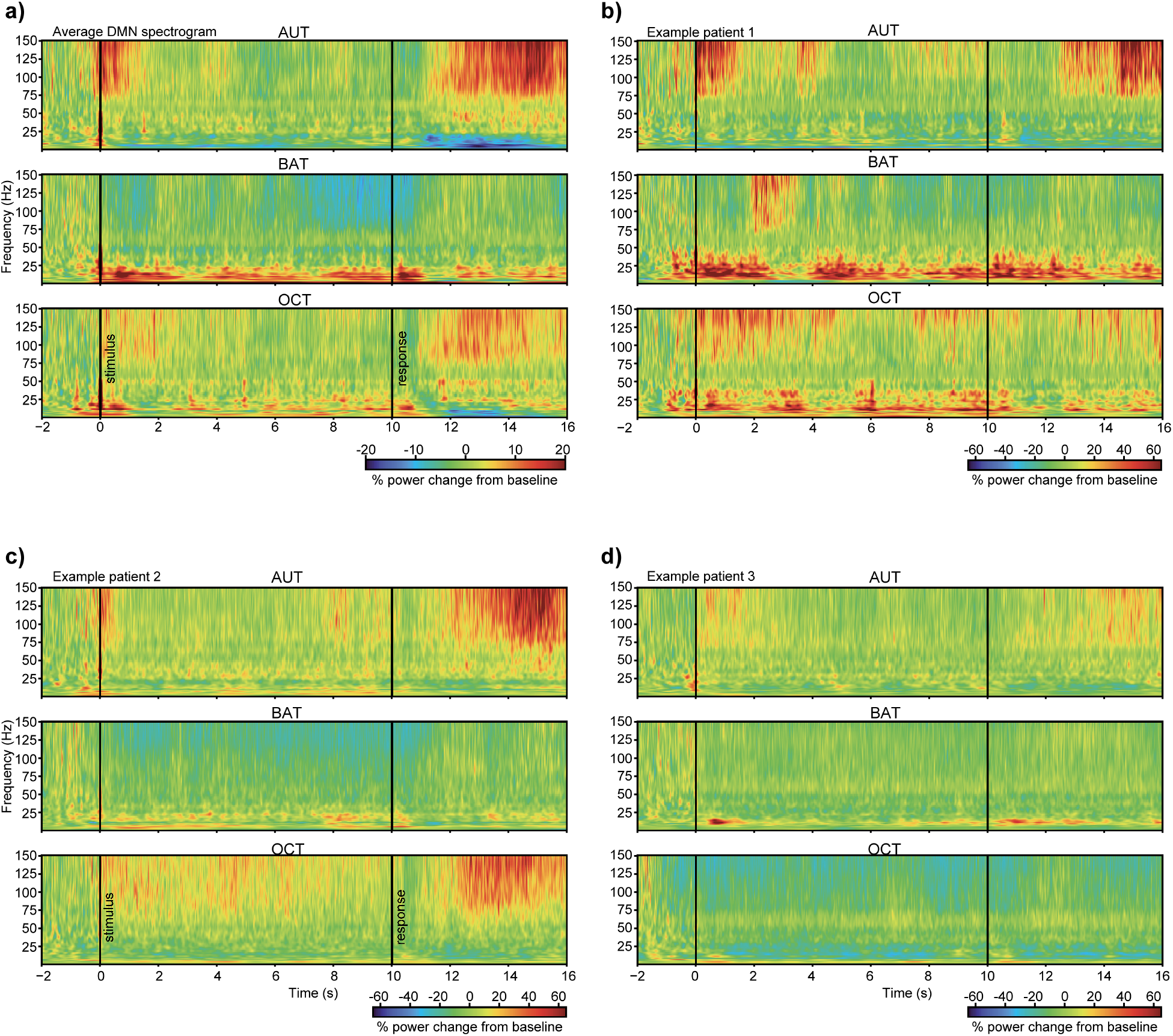
Average DMN spectrograms for the alternate uses task (AUT; top), basic arithmetic task (BAT; middle), and object characterization task (OCT; bottom). Each trial’s stimulus onset occurs at time 0. The data plotted before time 0 is the pre-block interval (the two seconds preceding the first trial of a block). The interval ranging from −2 through −1 seconds is averaged and used as a baseline for baseline correction. **a)** Spectrogram for all DMN channels averaged across patients (n=8). **b-d)** Spectrograms averaged across all DMN channels in each subject, for four example patients. Increased LFP power appears red/orange. Regions in which electrodes are located are as follows: **b)** UIC202307: 4 left middle temporal gyrus, 4 right middle temporal gyrus, 2 right medial frontal cortex. **c)** UIC202314: 7 left superior frontal gyrus (2 medial part), 1 left posterior orbital gyrus, 2 left lateral orbital gyrus, 2 left inferior frontal gyrus, 5 left middle frontal gyrus, 7 right middle temporal gyrus, 1 right inferior temporal gyrus. **d)** UIC202308: 2 left superior frontal gyrus, 1 left parahippocampal gyrus, 11 left middle temporal gyrus, 2 left superior temporal gyrus, 1 left posterior orbital gyrus, 1 left lateral orbital gyrus, 1 left inferior frontal gyrus, 3 right middle temporal gyrus.

**Figure 3.**
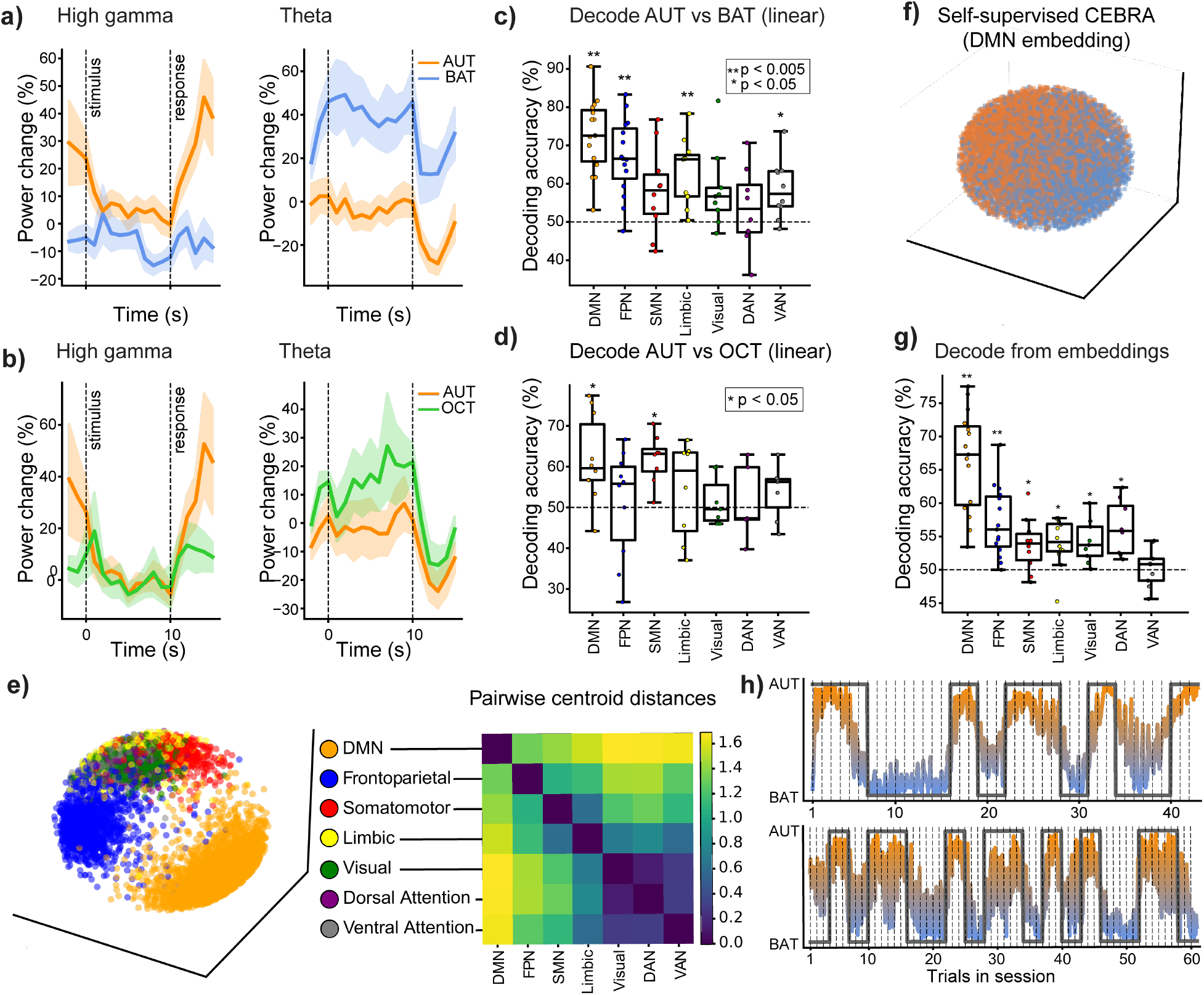
Default mode network representations distinguish creativity from control trials. **a)** Percent change from baseline of LFP power in high gamma (70-140Hz, left) and theta (4-8Hz, right) frequency bands over the time course of an average AUT (orange) and BAT (blue) trial. Baseline was calculated as the average of all activity during intervals 2 – 1 seconds before each block of trials. Difference between AUT and BAT before stimulus onset (time 0) is due to bleed-over from the previous trial (see Supp. Fig. 2). Shading indicates standard error. n=15 subjects. **b)** Same as (a), but comparing AUT (orange) and OCT (green). n=10 subjects. **c)** AUT vs. BAT decoding performance of linear SVM for each network. n=15 subjects. DMN: *p*=1.9×10-6; FPN: *p*=2.8×10-5; SMN: *p*=0.055; limbic: *p*=0.00055; visual: *p*=0.055; DAN: *p*=0.36; VAN: *p*=0.027 (two-sided 1-sample t-tests; FDR corrected for multiple comparisons). **d)** AUT vs. OCT decoding performance. n=10 subjects. DMN: *p*=0.022; SMN: *p*=0.006; FPN: *p*=0.827; Limbic: *p*=0.589; Visual: *p*=0.771; DAN: *p*=0.827; VAN: *p*=0.436; two-sided 1-sample *t*-test; FDR corrected. **e)** Dimensionality reduction using CEBRA trained to separate activity of different networks. Left: latent embeddings reveal network clusters. Each dot is 1 recording contact at 1 timepoint in a trial (16 total points per contact, because 16-second trial duration). Colors correspond to different networks. Right: pairwise centroid distances quantify distance in latent space between each pair of networks. Lighter (yellow) colors indicate greater difference; darker (blue) colors signify similarity between networks. Note that DMN is on average most different from the other networks, while visual network, DAN, and VAN are very similar to each other. **f)** Latent embeddings (for all AUT and BAT trials; n=15 subjects) generated by self-supervised CEBRA models based on DMN time series data (time-contrastive training; no behavior labels). Each dot is 1 timepoint. Embeddings are color-coded by trial type: AUT=orange; BAT=blue. **g)** KNN decoding of AUT vs BAT trial type from embeddings in (f) and Supp. Fig. 5, holding out 1 patient at a time when training CEBRA models, for cross-validation. Each dot is 1 patient. DMN: *p*=5.12×10^-2^; FPN: *p*=0.0009; SMN: *p*=0.02; limbic: *p*=0.006; visual: *p*=0.006; DAN: *p*=0.006; VAN: *p*=0.76; two-sided 1-sample *t*-test; FDR corrected. **h)** Projection of DMN embedding for two example patients onto a cognitive state coding axis defined by linear regression. Colored trace is the projection value; grey trace is the true trial type for each trial. Vertical dashed lines mark trial onsets. Abbreviations: default mode (DMN), frontoparietal (FPN), somatomotor (SMN), dorsal attention (DAN), and ventral attention (VAN) networks. In all box plots, center lines indicate medians; box boundaries indicate the 25th and 75th percentiles; whiskers extend to the most extreme values within 1.5 times the interquartile range.

Meanwhile, the control arithmetic task was characterized by decreased high frequency and increased low frequency power in DMN throughout the stimulus period. At the end of the stimulus, when AUT theta returned to baseline, BAT theta remained elevated. Upon entering the response window, theta power declined in both trial types, nearing baseline levels for BAT and falling below baseline for AUT. To reproduce the canonical DMN high gamma suppression during arithmetic, we tested the first 5 seconds after stimulus onset using a patient-level cluster-based permutation test of BAT high gamma power (in the first trial of each block) relative to a pre-task 4-minute resting baseline. In order to examine sub-second dynamics, we used a 0.1 second window size (as opposed to 1 second time bins in Figure 3a and the remaining analyses). This analysis revealed significant negative clusters at 0.3 – 0.8 s, 1.3 – 1.9 s, 2.0 – 3.5 s, and 4.3 – 5. Thus, BAT elicited significant early DMN suppression within the first second after stimulus onset, with a cluster from 0.3 – 0.8 s showing a mean −25.25% change from resting state (cluster-corrected *p*=0.0193), in line with previously reported timescales^27^.

We then quantified the difference between trial types. Across the duration of the entire trial (all stimulus and response time bins), high gamma power was overall greater for AUT than BAT (*t*_(224)_=8.27; *p*=1.22×10^-2^). When considering just the stimulus period (averaged across the window for each participant), we saw a significant difference between AUT and BAT theta power (*t*_(14)_=-3.79; *p*=0.002; two-sided paired *t*-tests) and high gamma power (*t*_(14)_=2.28; *p*=0.039), with BAT showing relatively higher theta and lower high gamma activity. Using cluster-based permutation tests to compare AUT vs. BAT high gamma power within the 10 second stimulus window, we found significant clusters at 0 – 2 s (AUT > BAT; cluster-corrected *p*=0.047) and 7 – 10 s (*p*=0.034) after stimulus onset. We also saw a significant difference in the 5 s response window, with a significant cluster at 11 – 15 s (*p*=0.0011).

In sum, since high frequency broadband activity reflects local neuronal population firing^28–32^ (while low-frequency LFP activity corresponds to network disengagement), these results suggest that DMN is disengaged during arithmetic and recruited during divergent thinking, replicating and expanding our previous findings on DMN power changes^19^. This relative disengagement also aligns with canonical observations of relatively reduced DMN high gamma during attention-demanding tasks^19,26,27^.

When comparing AUT vs. OCT (Fig. 3b), we find AUT characterized by greater response-period high frequency activity in DMN than OCT; OCT-related activity is intermediate between AUT and BAT. Low frequency (theta) power shows a similar pattern for OCT as BAT: theta power increases for OCT during the stimulus period (similar to BAT), while it decreases for AUT. During the response window, when BAT theta returns to baseline, OCT theta power drops below baseline, but does not exhibit as strong a decrease as does AUT theta. While AUT vs. BAT high gamma power was significantly different for multiple networks (all except Limbic), only DMN and VAN showed a significant difference between AUT and OCT high gamma (DMN: *t*_(159)_*=*2.85, *p*=0.005; VAN: *t*_(111)_*=*2.75, *p*=0.007; two-sided paired *t-*test comparing activity across all stimulus and response time points; FDR-corrected for multiple comparisons) with AUT showing stronger high gamma activity than OCT. Other networks either did not display a significant difference (FPN: *t*_(159)_*=*1.82, *p*=0.071; DAN: *t*_(79)_*=*-0.97, *p*=0.335; SMN: *t*_(127)_*=*0.59, *p*=0.558; Limbic: *t*_(127)_*=*-1.13, *p*=0.259) or showed stronger high gamma activity for OCT (Visual: *t*_(111)_*=-*3.54, *p*=0.0006), perhaps because describing an object’s physical characteristics required visualizing the object). Note that *t*-tests (averaged across time bins) did not reach significance for DMN. However, linear mixed effects models (with subject identity as a random effect) revealed significant differences between AUT and OCT: AUT exhibited significantly increased high gamma power compared to OCT during the response window (*p*=0.020); theta power was greater for AUT for the full trial and during both stimulus and response windows (full trial: *p*=1×10^-2^; early stimulus window (0-5 seconds): *p*=5.07×10^-2^; late stimulus (5-10 seconds): *p*=8.2×10^-2^; response: *p=*9×10^-2^).

In light of literature reporting a relationship between cognitive function and neural variability^33,34^, we also examined whether any differences in variability in high frequency power existed in the DMN between stimulus and response periods of the AUT, as well as between AUT and BAT trials. Though neither of the comparisons were significant, we found a slight trend toward larger trial-to-trial variability in DMN high frequency power for the creative condition compared to the arithmetic condition (*t*=1.50, *p*=0.16, two-sided paired *t*-test), as well as slightly higher response-period variability than stimulus-period variability within the AUT condition (*t*=1.73; *p*=0.11)—likely because high frequency activity in AUT is strongest during the response period (see Fig. 3a and our prior work^19^).

To further explore the neural patterns related to creative thinking, we applied machine learning methods to investigate how network activity encodes cognitive states (creative vs. arithmetic-related modes of thought, corresponding to AUT vs. BAT trial types). Using a linear support vector machine (SVM) classifier to decode cognitive state from the electrophysiology data, we were able to successfully distinguish trials involving divergent thinking vs. arithmetic on a trial-by-trial basis from DMN (72.98% decoding accuracy; Supp. Fig. 2a,b) and several other cortical networks—specifically the frontoparietal, limbic, and ventral attention networks; somatomotor, visual, and dorsal attention networks were *not* significantly predictive of cognitive state (Fig. 3c; Supp. Fig. 2a; DMN: *t*_(14)_= 9.2, *p*=1.9×10^-2^; FPN: *t*_(14)_=6.8, *p*=2.8×10^-2^; SMN: *t*_(9)_= 2.3, *p*=0.055; limbic: *t*_(11)_=5.3, *p*=0.00055; visual: *t*_(8)_=2.5, *p*=0.055; DAN: *t*_(7)_=0.97, *p*=0.36; VAN: *t*_(8)_=3.1, *p*=0.027; two-sided 1-sample *t*-tests; FDR corrected for multiple comparisons). To control for possible contamination from speech, we repeated this analysis by decoding only from the timepoints within stimulus window (during which subjects were silent). Decoding performance within the stimulus window was similar to that of the entire trial, except for the visual network, for which decoding performance crossed the significance threshold (DMN: *t*_(14)_=7.6, *p*=1.8×10^-2^; FPN: *t*_(14)_=4.7, *p*=0.0013; SMN: *t*_(9)_=1.0, *p*=0.38; limbic: *t*_(11)_=4.7, *p*=0.0015; visual: *t*_(8)_=3.8, *p*=0.0091; DAN: *t*_(7)_=0.66, *p*=0.53; VAN: *t*_(8)_*=*2.6, *p*=0.041).

Linear machine learning approaches also revealed a dissociation between AUT and OCT (Fig. 3d): cross-validated linear decoding (linear discriminant analysis) showed above-chance discrimination between AUT and OCT trials in DMN (61.83% accuracy, FDR-corrected *p*=0.022) and SMN (61.83%, *p*=0.006, but no other networks (FPN: 50.95%, *p*=0.827; Limbic: 54.30%, *p*=0.589; Visual: 51.45%, *p*=0.771; DAN: 51.37%, *p*=0.827; VAN: 53.78%, *p*=0.436; Fig. 3d).

The accurate cognitive state decoding suggests that divergent thinking depends on a unique network-wide pattern of neural activity and that distinct brain states underlie various modes of thought. Note that decoding analyses were performed separately within each participant and that group-level results reflect statistical aggregation of decoding performance across individuals rather than a single decoder trained on pooled neural data. Thus, cognitive state decoding results should be interpreted as evidence that distributed network-level patterns reliably differentiate cognitive states within individuals, rather than as a claim of a localized or invariant neural representation shared across individuals.

Although multiple brain networks distinguished creativity from arithmetic, DMN was the most predictive: classifiers trained on DMN activity significantly outperformed those based on somatomotor, limbic, visual, dorsal attention, and ventral attention networks (Fig. 3c; Supp. Fig. 2c; compared to SMN: *p*=0.0085; limbic: *p*=0.012; visual: *p*=0.012; DAN: *p*=0.0083; VAN: *p*=0.0085; two-sided Mann Whitney U test). DMN-based decoding also trended higher than frontoparietal-based decoding, though the difference was not significant (*p*=0.30). Note that variations in number of contacts between networks do not explain this difference: we observed no correlation between number of contacts and decoding performance in any given network (DMN: 8-29 contacts, *p*=0.85; FPN: 2-17 contacts, *p*=0.12; SMN: 1-7 contacts, *p*=0.80; limbic: 1-9 contacts, *p*=0.85; visual: 1-16 contacts, *p*=0.85; DAN: 2-9 contacts, p=0.85; VAN: 1-9 contacts, *p*=0.40; two-sided Pearson correlation). To further ensure that differences in electrode coverage did not skew our results, we reran the cognitive state decoding analysis for DMN and DAN only including patients with electrodes in both networks, and subsampling contacts so that the number of contacts matched between networks. For instance, for a patient with 5 DAN contacts but 10 DMN contacts, we randomly sampled 5 DMN contacts and performed the decoding analysis using just those contacts. When recomputing results for DMN using only the same number of contacts that were in DAN for each patient, we still found significantly above chance decoding performance (*p*=0.0012). Meanwhile DAN was not able to distinguish trial types (*p*=0.32), with DMN significantly outperforming DAN (*p*=0.038). (For contact-level analysis controlling for limited DAN coverage, see Supplementary Materials.)

After decoding cognitive state from network LFP activity, we inspected the classifier’s coefficient weights in order to identify which DMN subregions were most predictive of cognitive state: the features with the largest weights contribute the most to the classification and would indicate possible neurophysiological correlates for the decoding results. In 7 of the 15 datasets, the most predictive region was middle temporal gyrus (5 left; 2 right hemisphere). Superior frontal gyrus was most predictive in 3 datasets (2 left; 1 right), and superior temporal gyrus (left), inferior temporal gyrus (right), angular gyrus (right), posterior cingulate gyrus (right), and hippocampus (left) were each most predictive in 1 dataset.

To further interrogate the difference between cortical networks’ representations of cognitive state, we applied CEBRA, the recently developed nonlinear neural network encoder and dimensionality reduction tool^35^. Clustering network activity during only the AUT trials revealed a separation between DMN and other networks (Fig. 3e; Supp. Fig. 3), supporting the hypothesis that DMN has a unique role in creative thinking.

To corroborate the SVM decoding result and to visualize the separation between creative and arithmetic-related cognitive states, we employed CEBRA to extract the latent space that describes each dataset. We trained separate CEBRA models on the time series data for each patient, using trial type (AUT vs. BAT) as a supervisory signal (i.e., every timepoint for every trial was labeled as belonging to an AUT or BAT trial). We then used the model to transform the data, generating embeddings in a low-dimensional latent space. Importantly, to ensure proper cross-validation, we trained each model only on subsets of the trials (training set) and performed the same transformation on the training data and the held-out test data. When aligned between patients, the resulting manifolds clearly separated creativity from arithmetic states (Supp. Fig. 2b, left); moreover, embeddings from a held-out validation dataset displayed a similar pattern (Supp. Fig. 2b, right). We quantified this separation using a K-nearest neighbors (KNN) classifier to predict cognitive state based on the embeddings corresponding to each test subset (the held-out data). Decoding performance (Supp. Fig. 2d) was similar to that produced by the SVM, implying that cognitive state is linearly-separable at the network level. Similarly, CEBRA-generated embeddings separated AUT and OCT trial types (Supp. Fig. 4), with above-chance KNN decoding performance from embeddings based on held-out data for DMN (59.18% decoding accuracy) and most other networks (DMN: *p*=0.0085; FPN: *p*=0.0085; SMN: *p*=0.0086; Limbic: *p*=0.051; Visual: *p*=0.0086; DAN: *p*=0.011; VAN: *p=*0.0085; two-tailed one-sample *t*-test; FDR corrected).

**Figure 4.**
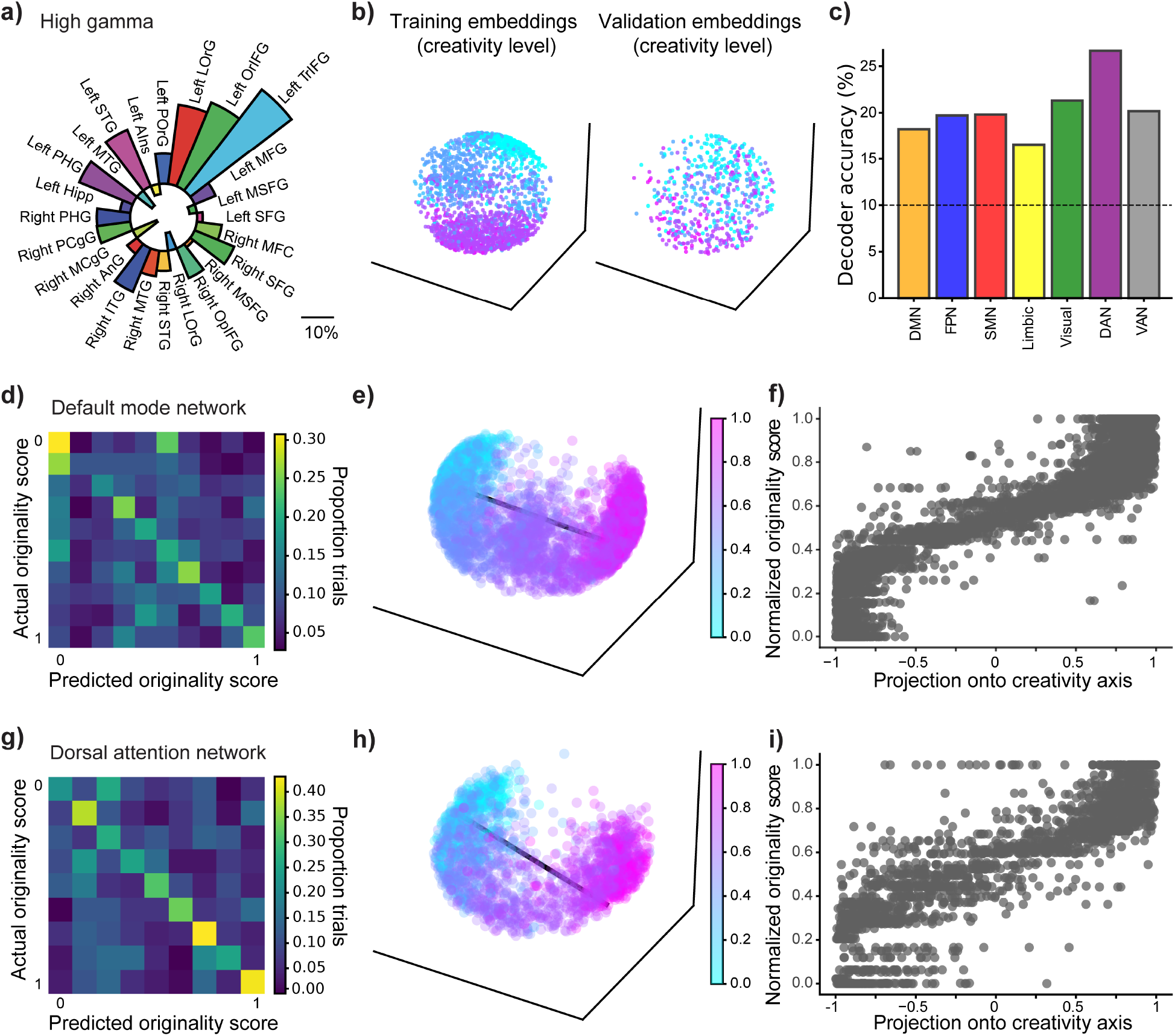
Dorsal attention network activity predicts performance on creativity trials through nonlinear representations. **a)** Difference in DMN network states between high vs low creativity: average high gamma LFP power (quantified as % change from baseline) of the ⅓ least creative trials is subtracted from the power of the ⅓ most creative trials. Outward pointing bars indicate a positive difference (i.e., greater power for high-originality trials); inward pointing bars correspond to negative difference. Scale-bar indicates magnitude of the difference. n=268 recording sites. **b)** Example of low-dimensional embeddings from the training and testing set of 1 cross-validation fold. Left: embeddings generated by CEBRA models trained on ¾ of the data for each patient to distinguish creativity levels. Right: Held out data transformed into the same latent space. Higher originality scores are pink; lower scores are blue. Each dot is 1 timepoint. **c)** Creativity decoding performance for each network based on the latent embeddings (average across 4 folds). Originality was discretized into 10 bins, so chance performance is 10%. **d)** Confusion matrix showing decoding performance based on DMN embeddings. Color indicates the proportion of trials for which a creativity score on the y-axis is predicted to be the value on the x-axis. **e)** Latent embeddings for DMN based on a CEBRA model trained on all data from all subjects. Black line is the coding axis determined by linear regression. Coding axis goodness-of-fit: *R^2^*=0.86. **f)** Projection of all datapoints of the latent embedding onto the coding axis shown in (e) (x-axis) compared to true normalized originality scores (y-axis). Y-axis for (f) is the same as the colorbar axis for (e). Pearson correlation *r*=0.93. **g)** same as (d), but for dorsal attention network. **h)** same as (e), but for dorsal attention network. *R^2^*=0.73. **i)** same as (f), but for dorsal attention network. *r*=0.86.

Our ability to decode trial-by-trial cognitive state from network activity, combined with the correlational analysis of LFP power that showed DMN engagement during AUT, reveals strong network-level representations of cognitive state, particularly in DMN.

### Cognitive state shapes the fundamental structure of DMN activity

In our decoding approach above, the CEBRA models were able to distinguish creativity vs. arithmetic trials when specifically trained to do so using trial type labels (AUT vs. BAT) supplied to the model. However, if cognitive state is a fundamental property of brain network activity patterns, it might be possible to predict trial type without providing the labels to the model. Hence, we asked whether an unsupervised or self-supervised model could still discriminate between creativity and arithmetic cognitive states.

To this end, we trained self-supervised CEBRA models using only time as a label, and no information about cognitive state, on the full dataset for each patient. To visualize the patterns related to cognitive state for each cortical network, we aligned across subjects and color-coded the latent space by trial type (Fig 3f; Supp. Fig. 5), revealing a separation between creativity and arithmetic trials that was qualitatively strongest in DMN. Subsequently, we attempted to predict cognitive state from these embeddings. Holding out one patient at a time, we again fit a KNN classifier to the remaining embeddings (training set) and asked it to classify trials for the held-out subject. Decoding performance was lower compared to that of the supervised models, but still significantly above chance for all networks except ventral attention network (Fig. 3g; median decoding performance for DMN: 67.29%, *t*=7.13, *p*=5.12×10^-6^; FPN: 56.04%, *t*=4.61, *p*=0.0009; SMN: 53.95%, *t*=2.93, *p*=0.02; limbic: 54.17%, *t*=3.44, *p*=0.006; visual: 52.72%, *t*=3.77, *p*=0.006; DAN: 55.83%, *t*=3.95, *p*=0.006; VAN: 50.83%, *t*=0.73, *p*=0.76; two-sided 1-sample *t*-test; FDR corrected). Furthermore, decoding performance was strongest for DMN (compared to FPN: *p*=0.003; SMN: *p*=0.0009; limbic: *p*=0.0005; visual: *p*=0.001; DAN: *p*=0.003; VAN: *p*=0.0005; two-sided Mann Whitney U test; FDR corrected).

**Figure 5.**
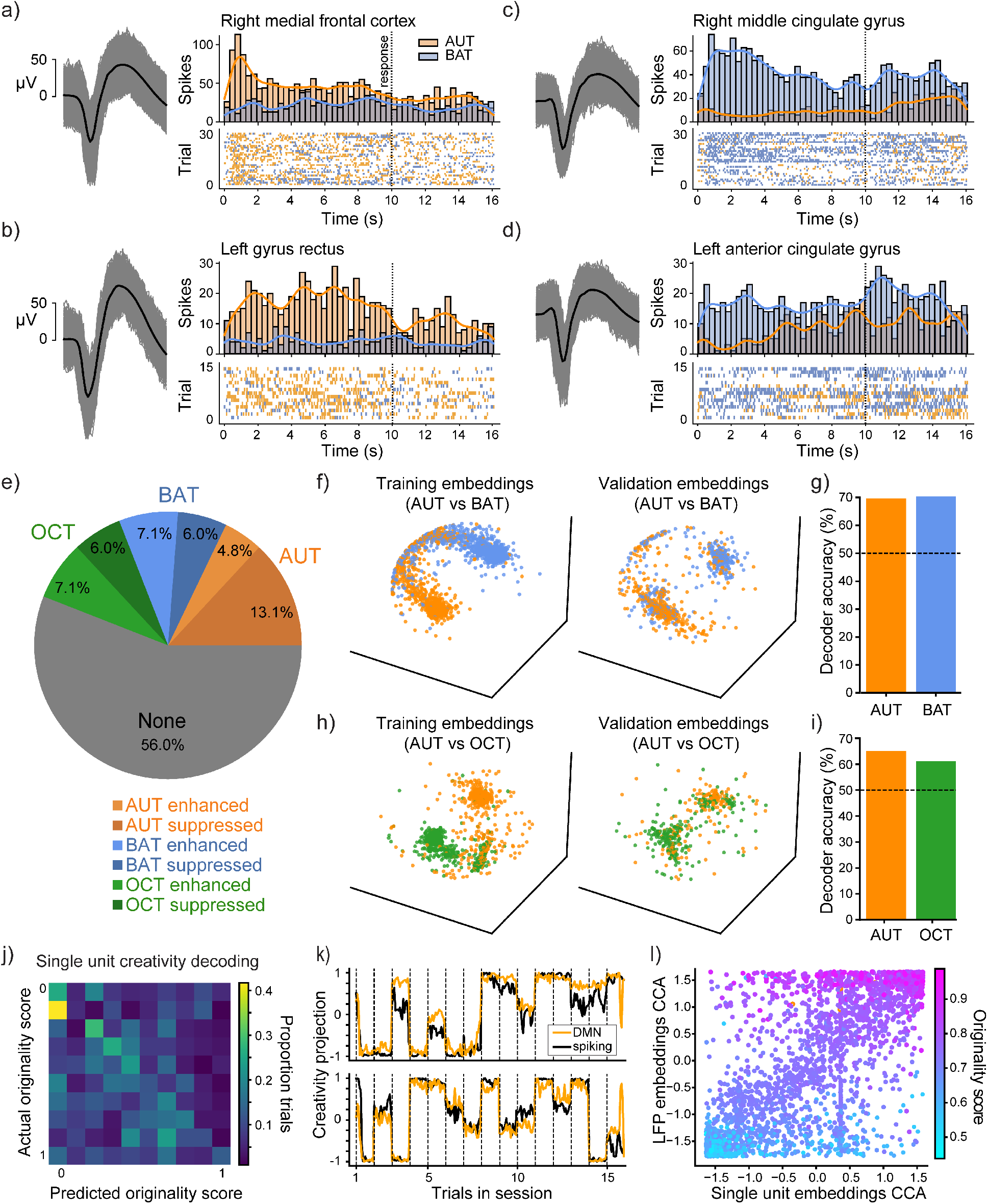
Trial type and creativity level are encoded at a single unit level. **a-d)** Four example units, recorded from (a) medial frontal cortex, (b) gyrus rectus, (c) middle cingulate gyrus, and (d) anterior cingulate gyrus. For each panel, Left: All waveforms of the identified unit. Light grey lines are individual waveforms; thicker black line is average. Right: PSTH (top) and raster plot (bottom) showing each cell’s responses to AUT (orange) and BAT (blue). PSTH shows total spike count for each 1 second time bin. For raster plot, each row is a trial, and each mark is a spike. Stimulus onset is at time 0. **e)** Proportions of cells identified as having a trial type preference (either selectively enhanced or suppressed in response to AUT, BAT, or OCT. n=85 cells for the 6 subjects with microelectrodes who completed all 3 tasks. FDR corrected. **f)** Separation of AUT vs. BAT trials (n=10 subjects; 127 cells). Example of latent embeddings from the training and testing set of 1 cross-validation fold. Left: embeddings generated by CEBRA models trained on ¾ of the single unit data for each patient to distinguish trial type. Right: Held out data transformed into the same latent space. **g)** Decoding AUT vs. BAT trial type based on latent embeddings, for AUT and BAT (n=10 patients). **h)** Same as (f) but separating AUT vs. OCT. **i)** Same as g, but decoding AUT. Vs. BAT (n=6 patients). **j)** Confusion matrix showing creativity level decoding performance based on single unit embeddings. Color indicates the proportion of trials for which a creativity score on the y-axis is predicted to be the value on the x-axis. *p*=0.007 (one-sided 1-sample *t*-test). **k)** Projection of single unit (black) and DMN (orange) data onto the creativity coding axis, for 2 representative patients. Vertical dashed lines indicate start of each trial. Y-axis is the projected creativity value. **l)** CCA between latent embeddings based on DMN and single unit data. Plot shows the first CCA component for the two data types (*r*=0.73). Color gradient indicates creativity value (*r*=0.88 for DMN and *r*=0.70 for single units).

Moreover, if we reduce the data even further to a single dimension along the coding axis (by projecting the data onto an axis defined by the weights of a linear regression model; see Methods), we find that its moment-by-moment trajectory tracks with the ground truth regarding creative vs. mathematical cognitive mode (Fig. 3h).

We remind the reader that this predictive latent space was generated by a self-supervised model, which had no knowledge of trial type information or behavior. Consequently, we can conclude that the representations of cognitive state are embedded in the intrinsic geometry of network activity, especially in DMN.

### Nonlinear network representations encode creativity level

We uncovered a relationship between neural activity patterns and creativity levels, as measured by the semantic distance between the prompt and the response for each trial. Specifically, we observed increased broadband high-frequency activity (high gamma; Fig. 4a) across DMN nodes in high-creative compared to low-creative states during the stimulus period of the AUT (*t*=1.78; *p*=0.038; one-sided paired *t*-test). The frontoparietal network also showed this pattern, but none of the other networks did (FPN: *p*=0.009; SMN: *p*=0.142; limbic: *p*=0.538; visual: *p*=0.296; DAN: *p*=0.356; VAN: *p=*0.158; one-sided paired *t*-tests). DMN low frequency (theta) power was lower on average for highly creative responses (Supp. Fig. 6a), but the difference did not reach the significance threshold (*p*=0.072). This finding of increased high gamma power in the most creative trials reveals state-specific DMN recruitment during highly creative states.

However, originality is not easily divided into two classes (high vs. low originality); rather, subjects’ responses comprise a distribution of semantic distance scores (Fig. 1b). Thus, creativity should be considered a continuous or multi-class metric. While a linear SVM was sufficient to decode creative vs. mathematical cognitive state with high accuracy, we were not able to decode levels of creativity with this methodology. Neither 2-class (high vs. low creativity) nor multi-class (creativity scores discretized into 10 bins) SVMs were able to accurately decode creativity scores, suggesting that any creativity representations, if present, are likely encoded nonlinearly.

Indeed, using a nonlinear dimensionality reduction approach through CEBRA, we were able to identify creativity level-related patterns in latent embeddings of the neural activity. To use this tool for decoding analyses, we trained a CEBRA model on subsets of LFP data (filtered into discrete frequency bands) from each subject, using discretized (binned and normalized) creativity score as a supervisory signal, and then applied the same transformation to the training set and held-out data to generate low dimensional embeddings (Fig. 4b). We then used the embeddings from the training set to decode discretized creativity level. We observed above-chance decoding performance for all sampled networks, while models trained on data with shuffled labels performed at chance level (Fig. 4c, Supp. Fig. 6b). However, unlike the cognitive state representations, for which DMN was the most predictive network, creativity level encoding was strongest in the dorsal attention network (Fig. 4c,d,g), with the dorsal attention network being significantly more predictive than DMN (p=0.02; two-sided Mann Whitney U test). The same pattern was observed when including timepoints from the entire trial as when only including the stimulus period; in fact, the stimulus-only embeddings yielded slightly higher decoding performance across all networks (Supp. Fig. 6b).

Given our ability to decode creativity level from neural activity, we asked whether the representations could be mapped onto a single dimension serving as a creativity axis (akin to coding axes identified for other cognitive or behavioral properties^36–38^). This axis can reveal moment-by-moment fluctuations in cognitive state, something that cannot be gleaned from univariate power or linear decoding. We generated new latent embeddings (Fig. 4e,h; Supp. Fig. 7), this time including data from all participants in one model for each brain network (treated by CEBRA as a pseudo-subject). In this analysis, continuous originality scores served as the labels for supervised learning. We then defined a coding axis via linear regression across embeddings (Fig. 4e,h; DMN: *R^2^*=0.77; DAN: *R^2^*=0.66). To confirm that this axis robustly described creativity performance, we projected the latent manifold onto this axis, producing a “creativity projection” value of the neural activity at each timepoint in each session. This projection correlated well with true originality scores (Fig. 4f,i; DMN *r*=0.87; DAN *r*=0.79 Pearson correlation), corroborating the presence of a creativity axis in the human brain while providing an instantaneous readout of a patient’s ongoing level of creativity.

Note that in this case, the relationship between DMN and creativity level appears stronger than that of dorsal attention network, an observation that holds even if we use normalized creativity values (see Methods for scores) and when we only compare data from the same subset of subjects (DMN subset: *R^2^*=0.75, *r*=0.87; DAN subset: *R^2^*=0.66, *r*=0.79). This dichotomy—better decoding from dorsal attention network but a more explanatory coding axis in DMN—could arise due the difference in the form of the supervisory signal: we used binned labels for classification analysis, but continuous labels to discern a coding axis. While both networks’ representations of creativity are nonlinear, having already gone through a nonlinear dimensionality reduction, we hypothesized that the dorsal attention network may be encoding creativity with a higher degree of nonlinearity, making it more suitable for discrete classification but less easily described by a linear regression model.

To investigate the nonlinearity of creativity coding, we fit polynomial regression models on the embeddings and normalized continuous originality scores. Compared to a linear model, a quadratic polynomial model described both networks’ representations of creativity better than the linear models did, with DMN models showing a more pronounced improvement. As we further increased the degree of the polynomial of the DMN models, the goodness of fit plateaued (i.e., greatest increase was between linear and quadratic). In contrast, DAN models showed the opposite pattern, with only small improvements in goodness of fit at first, and then increasingly larger improvements as the polynomial degree grew, overtaking the DMN model after a polynomial degree of 8 (Supp. Fig. 8a). This finding confirms the higher nonlinearity of the DAN creativity representation.

Finally, we considered the dimensionality of the representations. Simply performing extreme dimensionality reduction (using CEBRA to reduce the data down to a single latent dimension) could not identify a coding axis: cross-validated decoding performance based on a 1-dimensional latent space was at chance level for both networks (Supp. Fig. 8b). Thus, the coding axis we described above had to be embedded in a higher-dimensional latent space. Further, as the number of latent dimensions increases from 1 to 5, the difference between decoding based on DAN vs DMN widens (and then plateaus after 5 dimensions), suggesting that DAN representations of creativity are more high-dimensional.

Together, these findings expose brain-wide network-level creativity representations that are encoded along a nonlinear coding axis in a high-dimensional latent space. Further, these results hint at a role of the dorsal attention network in modulating the quality of individual subjects’ creative output through a highly nonlinear representation of creativity.

### Individual differences in originality correspond to network connectivity

So far, we have shown how creativity-related information is represented at the network level. However, we have not yet considered inter- and intra-network interactions—especially within and between DMN and FPN—which have been previously implicated in creative thought^15,39,40^. Consequently, we investigated phase synchronization within and between these two networks, in which we had substantial coverage across subjects. We measured phase coherence based on phase locking between contact pairs across DMN and FPN within the canonical frequency bands and quantified the relationship between maximum coherence during the stimulus period and creativity level. Mixed-effects models accounting for participant-level clustering did not identify a consistent trial-level within-subject relationship (DMN-FPN, *β*=-0.001, *p*=0.936; FPN, *β* = 0.002, *p*=0.877). By contrast, patient-level aggregate analyses suggested a between-subject association (Supp. Fig. 9, top): subjects with higher mean originality tended to show higher mean gamma (30-70 Hz) synchrony in FPN (*β*=1.07, *p*=0.014, *R²*=0.385), with a similar trend for DMN-FPN (*β*=0.342, *p*=0.063, *R²*=0.242). These exploratory results suggest that the strength of within-FPN (and possibly DMN-FPN) coupling reflects between-subject individual differences (rather than within-participant trial-level coupling). For theta (4-8 Hz), FPN theta coherence during the stimulus window shows a negative within-patient relationship with originality in the mixed-effects model (*β*=-0.061, *p*=0.041), while DMN and DMN-FPN do not show reliable associations; patient-level models showed no significant between-participant relationship (Supp. Fig. 9, bottom).

### Cognitive state is encoded at single unit level

Given the large-scale, network-level representations of cognitive state we uncovered, we next asked whether individual cells represent these distinct cognitive states. In a subset of patients (10), we recorded single-unit activity in addition to LFP responses. Indeed, we found cells in multiple brain regions that were selective for trial type (creativity vs. math cognitive states): some exhibited increased firing for AUT trials (Fig. 5a,b) while others preferred BAT trials (Fig. 5c,d). Of the 127 cells we recorded in the 10 patients who performed AUT and BAT, 41% significantly encoded creative vs. arithmetical cognitive state (two-sided paired *t*-tests; FDR corrected). Of these selective cells, 43.9% were more responsive for AUT trials. For the 6 subjects (85 cells) who performed AUT, BAT, and OCT, we found 44% of cells with trial type preference (Fig. 5e; one-way ANOVA to determine cells with significant selectivity, FDR corrected; two-sided paired *t*-tests to determine which single trial type elicited significantly different response in the cell from the other two trial types; see Methods). 17.9% of cells showed AUT responses significantly different from both BAT and OCT responses; 13.1% of cells showed BAT-selectivity; 13.1 % of cells were selective for OCT. Selective cells had either enhanced or suppressed firing rates in response to one of the 3 trial types.

Selective cells were identified in the following regions: medial temporal lobe (N=47, 29.29% selective), ventromedial prefrontal cortex (N=37; 40.54% selective), dorsomedial prefrontal cortex (N=15; 53.33%), middle cingulate cortex (N=20; 45%), and posterior cingulate cortex (N=8; 50%).

Further, the neuronal population’s representations of different cognitive states were separable (Fig. 5f-i). We trained CEBRA models on the single-unit responses for each patient, either contrasting AUT vs. BAT trials (Fig. 5f; n=10 subjects) or contrasting AUT vs. OCT trials (Fig. 5h; n=6 subjects), and from the resulting embeddings, we were able to distinguish AUT vs BAT trials (Fig. 5g; *p*=0.0009; two-tailed one-sample *t*-test) and AUT vs. OCT trials (Fig. 5i; *p*=0.016). Thus, the single-neuron responses among the population of recorded cells reflect engagement in creative vs. other forms of thought.

### Single unit and network activity encode creativity similarly

Finally, we investigated the relationship between creativity representations in the network-level LFP activity and the single-unit spiking patterns. As with LFP data, we used CEBRA to identify the single unit representations of creativity for each patient (holding out data from CEBRA training with 5-fold cross-validation). We found that not only cognitive state but also creativity level was encoded in the responses of the recorded neuronal population. Classification performance on single unit responses was lower than on network-level LFP activity—which is to be expected given the low number of units (features on which the model is trained) in each patient. Nonetheless, creativity scores could still be decoded above chance from embeddings based on the single unit population (Fig. 5j; *p*=0.007).

We then computed the creativity coding axis using linear regression on the single unit spiking data. Then we projected the data onto this coding axis and compared the result to the LFP creativity projection. We observed close alignment between the two projections (Fig. 5k), suggesting shared representations between these two data sources.

To quantify this relationship further, we employed canonical correlation analysis (CCA) on the creativity representations (low-dimensional embeddings generated by CEBRA) corresponding to single unit and DMN LFP activity. We found a strong correlation between the LFP and single unit embeddings (Fig. 5l; first CCA component correlation between spiking embeddings and DMN embeddings: *r*=0.73; FPN: *r*=0.70; SMN: *r*=0.65; limbic: *r*=0.58; visual: *r*=0.68; DAN: *r*=0.69; and VAN: *r*=0.74), demonstrating that LFP and spiking activity share a dominant latent dimension in the creativity embedding space and thus revealing the close relationship between the single unit spiking and network-level representations of creativity.

Both LFP and single unit embeddings displayed high correlation between their first CCA component and creativity scores (*r*=0.88 for DMN and *r*=0.70 for single units). The higher correlation of LFP embeddings with creativity suggests that this representation describes more of the variance in creativity performance than do single unit embeddings, a difference that may arise from the more widespread coverage of DMN by LFP data compared to limited and local single unit recordings. Nevertheless, these results validate the LFP findings and reveal shared creativity representations between single neuron firing and large-scale brain networks.

## Discussion

Despite growing interest in the neuroscience of creativity, the neural geometries underlying the emergence of novel and diverse ideas remain mysterious. Our investigation of a neural basis for creativity unearthed widespread encoding of cognitive state and originality of creative thought. Indeed, we found that unique brain states distinguished creative thinking and arithmetic calculations, allowing us to disambiguate these cognitive states from the activity of several brain networks, including DMN, frontoparietal, limbic, and ventral attention networks. Of these networks, we identified DMN as the most linearly predictive of cognitive state. DMN also distinguished creativity from a semantic, autobiographical object characterization task.

Not only was DMN predictive of cognitive state, but also its activity structure was intrinsically organized to reflect this feature: a self-supervised dimensionality reduction approach revealed patterns of activity that distinguished creativity from mathematical thinking. This separation of cognitive state was much clearer in DMN than in any other network. Importantly, this finding is not inevitable: just because the trial type information can be extracted from DMN activity does not imply that this variable is a fundamental property of DMN activity, and we did not initially expect to be able to distinguish modes of thought using a model with no supervisory signal or behavioral information. The striking observation that a self-supervised model spontaneously separated creativity and arithmetic states indicates that cognitive state is inherent to the structure of DMN activity (i.e., intrinsically embedded in its representational geometry).

The self-supervised CEBRA results also indicate that representations align and generalize across subjects, regardless of electrode coverage or specific brain regions involved. In other words, creative ideation is not encoded at specific recording locations, but rather supported by flexible, population-level dynamics that can be instantiated through different anatomical substrates, but which preserve consistent large-scale network organization.

While the creative state (compared to math) was most strongly encoded by DMN, it was the dorsal attention network that contributed most to varying levels of creative output. This finding was especially surprising given that the dorsal attention network was the least predictive of cognitive state. Although the DAN results should be interpreted with caution given the limited DAN coverage in our dataset, we speculate that the DMN plays a role in gating creative thought, while the dorsal attention network is more involved in regulating levels of originality (as assessed by the alternate uses task), albeit with a highly nonlinear representation. Creative ideation is thought to involve two parallel processes: idea generation and evaluation. Both processes likely occur during our divergent thinking task, though our behavioral design does not dissociate between them. Due to its involvement in episodic memory and self-referential thinking^10,41,42^, DMN is believed to play a stronger role in idea generation, during which thoughts arise spontaneously or through associative thinking^43^, while networks related to executive control and goal-directed thought contribute more to idea evaluation^15^ However, our speculation regarding the gating vs. regulating aspects of creativity, though akin to generation and evaluation, was not based on a temporal theory of creative thought phases. Rather, we imagine near-simultaneous processes that generate and constrain creative thought in our short trial structure. In addition, self-referential processing within the DMN is likely not unitary and may reflect transient engagement of distinct but nested processes, as proposed in hierarchical models of the self^44^.

Accordingly, a gating DMN may help a participant think of ideas, but they may not necessarily be good ideas; the dorsal attention network may then contribute to fine-tuning associative thought to produce a final response, through allocation of attentional resources and prioritization of task-relevant information^7^. Since we only have access to the final spoken response, the potential DAN involvement in choosing and articulating that response could explain why DAN activity most strongly predicts the quality of the final output. Importantly, our dataset contains limited DAN coverage, so the DAN results should be considered exploratory and not as evidence of a fundamental mechanistic role of DAN in modulating creativity.

Interestingly, this hypothesized top-down role is often associated with the frontoparietal control network, rather than with the attentional system. However, since the dorsal attention network is believed to control goal-directed attention^45^, we considered the dorsal attention to play a distinct but complementary role in creativity^14,46^, potentially providing “automatic” constraints during both idea generation and evaluation^14,47^ in addition to possible top-down attentional control. Li et al. (2026) report that DAN shows stronger influence over other networks in highly creative individuals—possibly because creative thought depends on attentional maintenance of complex goal-related information as well as efficient switching between streams of thought^48^. It is also possible that the frontoparietal network mediates the dorsal attention activity patterns we observe or that coupling with FPN may link the roles of DMN and DAN^7,49^. Indeed, we found that the strength of FPN (and to a lesser degree, DMN-FPN) gamma-band coupling reflects a participant’s average creativity in the AUT. Although the interpretability of spectral connectivity results is complicated by idiosyncrasies in neural activity patterns across human epilepsy patients, by high variability across trials and time, and by limited access to internal decision-making processes, our exploratory phase coherence analysis suggests a possible modulatory role of FPN. Thus, we cannot speculate as to whether the control of creative thought originates in DAN or FPN—only that the information related to the originality of participants’ final responses is more strongly represented in the dorsal attention network.

This framework, with DMN gating divergent thinking, is also supported by previous research in which disruption of lateral DMN nodes through high-frequency stimulation decreased the number of ideas generated (a metric known as “creative fluency”) but did not alter originality scores^11^. Indeed, in the current study, the regions most predictive of cognitive state were predominantly localized in lateral DMN in 10 of the 15 patients (7 middle temporal gyrus, 1 superior temporal gyrus, 1 inferior temporal gyrus, and 1 angular gyrus), while dorsal and medial regions were more predictive in only 5 subjects (3 superior frontal gyrus, 1 posterior cingulate gyrus, and 1 hippocampus). Additionally, our more recent work^19^ revealed that DMN recruitment during AUT was driven by activation in lateral DMN nodes. Conversely, stimulating medial (rather than lateral) DMN nodes interfered with originality, but not fluency, suggesting that distinct anatomical-functional sub-specializations within the DMN influence quantity and quality of creative output.

Besides different anatomical targeting, task design variations may also explain the diversity of past studies’ results. Consider the task differences from the generation-evaluation framework: when given enough time to engage in mind-wandering and permission to verbalize each idea (as in our previous AUT experiments), idea generation is likely to be the dominant mode of thought^19^. In contrast, a more engaging task with a shorter stimulus window and instructions to only report a single response (as in the current task design) may increase reliance on the idea evaluation subprocess. Thus, distinct mechanisms likely drive divergent thought in our current paradigm, with greater emphasis on executive control and top-down decision making.

The present study utilized the AUT as a well-established divergent thinking task with inherent constraints—time-limited design, specific task instructions, and only one facet of creative cognition examined. These constraints facilitated the application of sophisticated computational analysis that yielded critical insights; however, they also result in limitations like the need for attention and cognitive effort (which may be absent during spontaneous creative thought). Although effort is inherently tied to creativity in our present design, effort alone cannot explain our observed results: for instance, existing literature demonstrates that increased cognitive effort is typically associated with DMN deactivation^50,51^, rather than increased DMN engagement, which we observed in AUT and in more original trials. However, future research will explicitly tackle this distinction by implementing behavioral paradigms that probe spontaneous (rather than instructed) creative ideation.

Besides the inter-network coding differences, we also discovered a brain-wide creativity-coding axis. The projection of creativity-related manifolds onto this axis (i.e., the moment-by-moment representation of creativity level) correlated well with true originality scores. Thus, despite the multidimensional geometry of the latent space (corresponding to even higher-dimensional neural data), a discrete coding axis can be isolated. Furthermore, this axis is preserved across scales, from single units to distributed networks, suggesting that creativity is a hierarchically-organized, multiscale process.

The creativity axis we identified enables the prediction and instantaneous readout of moment-by-moment creativity state as it evolves over the course of a trial. Although our analyses focus on creativity representations rather than explicit linguistic dynamics, the moment-by-moment fluctuations along the creativity-coding axis raise the possibility that these signals reflect rapid semantic processes involved in formulating a response. Although these processes cannot be explicitly resolved with the current task structure (brief responses; silent stimulus window), our findings align with recent work on the temporal dynamics of semantic cognition^52^. However, as mentioned previously, we do not speculate as to whether each timepoint’s creativity state belongs to any given “phase” of creative ideation (e.g., idea generation; evaluation).

When we projected all datapoints in each network’s latent space onto the creativity-coding axis, the correlation of the resulting projection with true creativity values was highest for DMN compared to the other networks, including dorsal attention network. Similarly, the goodness of fit of the linear regression model was higher for DMN than for dorsal attention network, meaning the creativity coding axis in DMN explained the data better. Yet, earlier, we discussed our observation that dorsal attention network, not DMN, was most predictive of creativity level. Though these analyses both delve into similar questions about network-level representations of creativity level, there is a key difference: the decoding analysis was based on embedding generated from binned creativity data, whereas the coding axis was inferred based on continuous originality scores. We speculate that the dorsal attention network’s representation of creativity is higher-dimensional and more nonlinear. This conjecture is supported by our observation of improved fit of curved coding axes generated by higher-polynomial regression models, and that increasing the degree of the polynomial has a stronger effect on the dorsal attention network than DMN. Thus, it was possible to recover a latent space from DMN that encodes creativity as a continuous spectrum along a near-linear coding axis spanning all creativity levels; meanwhile, the same approach revealed that the dorsal attention network’s representation is highly nonlinear. This result is consistent with our proposed framework in which the dorsal attention network regulates creativity through goal-directed attentional control of thought processes: unlike DMN’s simpler and stable relationship with creativity, the dorsal attention network generates a highly complex representation of creative states, which may fluctuate with task demands and cognitive engagement.

Finally, our examination of single-unit recordings reveals consistent geometries across scales, and points to a potential mechanism for creativity encoding at the single-cell level that may underlie network-wide creativity representations. The creativity coding axis we observed in network-wide LFP data was also found in single-unit spiking patterns, and the trajectories of the creativity-trained embeddings based on the LFP and single-unit data closely followed each other. Therefore, the way creativity is represented by the spiking of single units is largely conserved at the level of network activity. We found this relationship between spiking and network representations for each network sampled, likely because we considered all recorded units as a single population and did not split them according to anatomical or functional boundaries. Future work will collect a larger sample of single units and enable more fine-grained analysis. In addition to single unit recordings, stimulation experiments to manipulate network activity and synchrony will be important to clarify the role played by specific neuronal ensembles in generating creative thought.

In the meantime, we conclude that the default mode network maintains a stable representation of cognitive state, while high-dimensional nonlinear brain states across networks flexibly encode creativity level. Although cognitive state and creativity level can be decoded from activity brain-wide, the default mode network likely plays a key role in gating creative thought, and its activity patterns are uniquely shaped by creative ideation. In contrast, our results suggest that the dorsal attention network generates a complex, high-dimensional representation of creativity level that imposes context-dependent constraints and flexibly regulates the quality of creative output. Together, our findings serve to illuminate the neural landscape underlying creative thought.

## Methods

### Subjects

18 neurosurgical patients (Table 1) who were undergoing clinical monitoring for epilepsy participated in the study. All subjects provided informed consent prior to participation. Separate informed consent was obtained for implantation of microelectrodes and for participation in cognitive tasks. All experimental procedures were performed according to protocols approved by the University of Utah Institutional Review Board. 1 patient (UIC202420) was excluded from analysis due to experiment interruption for clinical care (thus leaving n=17 participants included in the paper). Only patients aged 18 and older participated in the experiments (mean age = 40.1 ± 11.2 years; range = 27 - 65). No patients with major neuroanatomical anomalies or cognitive deficits were included.

**Table 1:**
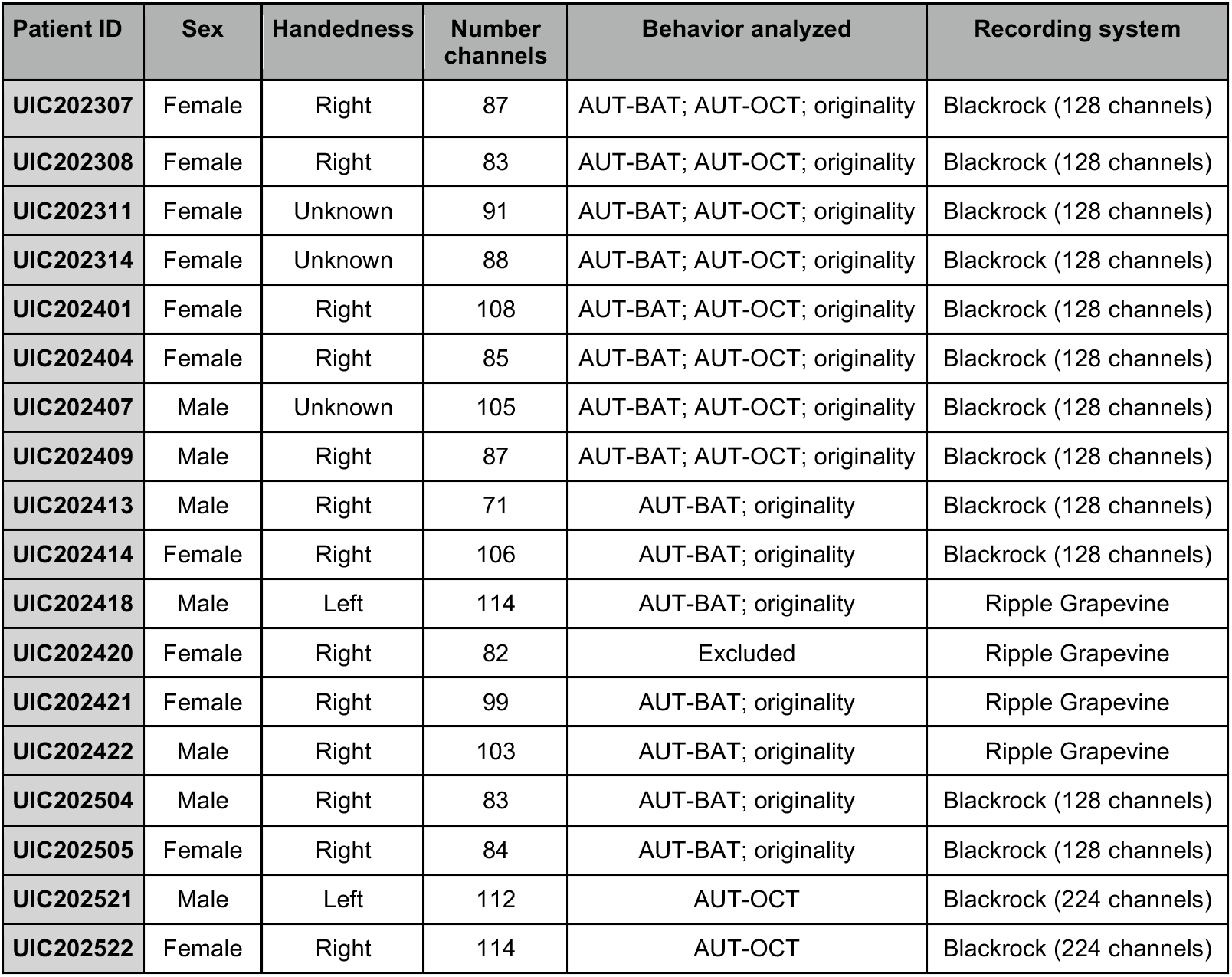
Participant demographics. Table shows demographics for included participants. Number of channels is the number of macrocontacts for each patient after removing noisy channels. UIC202420 was excluded from analysis due to clinical interruption.

| Patient ID | Sex | Handedness | Number channels | Behavior analyzed | Recording system |
|---|---|---|---|---|---|
| **UIC202307** | Female | Right | 87 | AUT-BAT; AUT-OCT; originality | Blackrock (128 channels) |
| **UIC202308** | Female | Right | 83 | AUT-BAT; AUT-OCT; originality | Blackrock (128 channels) |
| **UIC202311** | Female | Unknown | 91 | AUT-BAT; AUT-OCT; originality | Blackrock (128 channels) |
| **UIC202314** | Female | Unknown | 88 | AUT-BAT; AUT-OCT; originality | Blackrock (128 channels) |
| **UIC202401** | Female | Right | 108 | AUT-BAT; AUT-OCT; originality | Blackrock (128 channels) |
| **UIC202404** | Female | Right | 85 | AUT-BAT; AUT-OCT; originality | Blackrock (128 channels) |
| **UIC202407** | Male | Unknown | 105 | AUT-BAT; AUT-OCT; originality | Blackrock (128 channels) |
| **UIC202409** | Male | Right | 87 | AUT-BAT; AUT-OCT; originality | Blackrock (128 channels) |
| **UIC202413** | Male | Right | 71 | AUT-BAT; originality | Blackrock (128 channels) |
| **UIC202414** | Female | Right | 106 | AUT-BAT; originality | Blackrock (128 channels) |
| **UIC202418** | Male | Left | 114 | AUT-BAT; originality | Ripple Grapevine |
| **UIC202420** | Female | Right | 82 | Excluded | Ripple Grapevine |
| **UIC202421** | Female | Right | 99 | AUT-BAT; originality | Ripple Grapevine |
| **UIC202422** | Male | Right | 103 | AUT-BAT; originality | Ripple Grapevine |
| **UIC202504** | Male | Right | 83 | AUT-BAT; originality | Blackrock (128 channels) |
| **UIC202505** | Female | Right | 84 | AUT-BAT; originality | Blackrock (128 channels) |
| **UIC202521** | Male | Left | 112 | AUT-OCT | Blackrock (224 channels) |
| **UIC202522** | Female | Right | 114 | AUT-OCT | Blackrock (224 channels) |

### Electrophysiological recordings

Depth electrodes were implanted in patients’ brains as part of their epilepsy monitoring, and the location of the implants was dictated by clinical need. 0.8mm diameter DIXI SEEG electrodes were used, containing macrocontacts with 3.5mm spacing between contacts and 2mm contact length. For the patients who consented (n=10), 2-3 electrodes used were Behnke-Fried hybrid electrodes (1.38mm diameter), which contained both macrocontacts (5mm spacing; 1.57mm length) as well as 40um microwires at the end. These microwires splay out during implantation and provide recordings of single unit activity. All other electrode contacts recorded local field potentials (LFP).

Recording sites were classified as belonging to a particular region and brain network using the LeGui software package^53^, based on a post-operative CT scan and co-registered structural MRI and normalized to MNI space (warping from patient space to MNI space was required due to anatomical variations between subjects). Regions were identified using the Neuromorphometrics (NMM) atlas and networks were identified using commonly-accepted atlas-based network parcellation^11,15,19,54–60^ using the Yeo 7 network atlas^61^. White matter contacts were excluded from analysis. Only contacts identified as belonging to one of the 7 networks were included. Contacts outside of atlas-defined network boundaries were not labelled as belonging to any of the 7 networks and excluded from analysis. Contacts with unclear network or white/grey matter classification (e.g., with <0.1 probability of belonging to both grey and white matter) were also excluded. Contact classification by network, region, and grey/white matter was performed using LeGUI software. (See Davis et al., 2021 for detailed methodology on contact classification by network, region, and grey/white matter).

The total number of channels in each network, and the number of patients with coverage was as follows:

For the cohort of 15 subjects who completed the AUT-BAT contrast, the values were as follows: DMN: 268 channels across 15 subjects; FPN: 120 channels, 15 subjects; SMN: 40 channels, 10 subjects; Limbic: 45 channels, 12 subjects; Visual: 58 channels, 9 subjects; DAN: 37 channels, 8 subjects; VAN: 31 channels, 9 subjects. For the 10 subjects who completed the AUT-OCT contrast, the values were: DMN: 162 channels; 10 subjects; FPN: 72 channels, 10 subjects; SMN: 45 channels, 8 subjects; Limbic: 32 channels, 8 subjects; Visual: 48 channels, 7 subjects; DAN: 21 channels, 5 subjects; VAN: 23 channels, 7 subjects.

14 sessions were conducted using a Blackrock Microsystems recording system (128 channels (12 patients) or 224 channels (2 patients), 1KHz sampling rate); 3 sessions were recorded with a Ripple Grapevine system (128 channels, 2KHz sampling rate, downsampled to 1KHz for analysis). Single unit activity was recorded from 10 subjects (all 10 subjects performed AUT and BAT; 6 subjects also performed OCT).

### Behavior

#### Behavioral paradigm

15 of the 17 subjects performed the alternate uses task (AUT) and a basic arithmetic task (BAT) control; 10 of the 17 subjects performed the AUT and an Object Characterization Task (OCT) control (see Table 1). All three tasks had identical structure. AUT required participants to generate the most creative use of the presented item; BAT asked subjects to solve an arithmetic equation; OCT asked participants to report the most defining characteristic of the item, which could include the common (noncreative) use of the item.

Prior to initiating the task, the experimenter explained the instructions and task structure in detail, including examples of possible creative uses for an example prompt (“key”) to ensure participants understood the task. experimenter emphasized that creative responses should be unusual and non-obvious uses of the objects. Because the task required only a single verbal response per trial, participants were instructed to select and report the use they personally considered most creative if multiple ideas were generated. However, the brief trial structure (only 10 seconds to think of an answer) meant that participants were unlikely to think of multiple responses. Then, subjects were exposed to a “training round” to help them understand the task and to ensure that all subjects interpreted the instructions the same way. The training round also allowed the experimenter to provide feedback. The training round included AUT and the control task(s); once subjects were comfortable with task structure, the training ended and the real task began. The tasks were designed using psychtoolbox.

Each trial consisted of a 10 second stimulus window followed by a 5 second response window. Each block of trials contained 3 trials of one type. AUT and control (BAT; OCT) trials were randomly interleaved. At the beginning of each block, instructions were shown for the corresponding trial type. For each trial within a block, a prompt (cue word or arithmetic equation) was presented on a screen for 10 seconds, during which subjects were instructed to silently think of their answer. For AUT, the instructions and cue word were displayed in orange, for BAT, the font was blue, and for OCT, the font was green. After the 10 second stimulus window, the prompt text changed to white, signaling subjects to verbally respond by either stating a single "most creative" use for the object or providing the answer to the equation.

Each session included 15-30 trials of each type. AUT and OCT prompts were randomly selected from a list of 100 common items (e.g., spoon, newspaper, hanger) without repetition. For each BAT trial, subjects were prompted to add or subtract two random numbers between 0 and 12 (e.g., 8+5 or 4-12). We chose simple arithmetic to avoid the engagement of creative problem-solving that might be required for more complex mathematic reasoning.

#### Performance scoring

Participants’ responses were collected with a microphone, and transcribed offline. AUT performance was judged based on the originality of each response. Originality was quantified as the semantic distance between the prompt and the response, using the Open Creativity Scoring algorithm^62,63^, which is specifically optimized for scoring the AUT.

Although we previously used frequency-based metrics of originality^11^ (related to the rate of occurrence in the dataset), frequency-based metrics suffer due to their dependence on subject sample size (and heterogeneity), phrasing/vocabulary, and task constraints. Semantic distance, with its continuous and stable measure of conceptual novelty, is more robust to these variables, generalizing beyond the specific patient population. It also allows conceptually similar responses to be treated equivalently even when phrased differently (while frequency-based analysis can be “tricked” by lexical rather than conceptual novelty). Furthermore, a continuous within-subject spectrum of originality scores was critical for our trial-by-trial neural analysis, decoding, and the discovery of a creativity axis in the brain. Finally, our version of the task uses a large pool of possible prompts in order to reduce stimulus identity-specific confounds and improve generalizability, meaning each prompt had relatively few responses associated with it, so frequency-based analysis would be under-powered in our paradigm.

Notably, semantic distance may correlate with abstraction, increasing when responses move away from concrete, conventional uses toward more abstract or conceptually distant associations. This characteristic is not a limitation of the measure but rather reflects a core component of originality in divergent thinking tasks such as the AUT. Generating abstract or conceptually distant uses requires accessing less typical semantic associations, which is precisely what semantic distance is designed to capture. However, semantic distance does not prioritize abstraction, but rather the distance in semantic space relative to the object prompt. Abstract responses that remain closely related to the object’s common use receive lower scores, while concrete responses that are semantically distant receive high scores. Thus, semantic novelty, not abstraction and concreteness, is the defining dimension of this score. Consistent with prior work, we therefore use semantic distance as a standardized measure of originality^15,19,22–25^.

The Open Creativity Scoring algorithm calculates originality as follows: after removing stopwords (e.g., “the”, “and”, “to”), this natural language processing algorithm uses a semantic embedding model to convert both prompt and response to vectors in semantic space and calculate the cosine of the angle between the vectors. The larger the angle, the more dissimilar the response is from the prompt, and the greater the originality score.

### Analysis of LFP power

The Python MNE library^64^ was used to analyze LFP data. Data preprocessing included removing noisy channels, notch filtering to remove power line noise (60Hz and 120Hz harmonic), and common average referencing. Data was epoched to only include relevant time windows. Frequency bands were split as follows: theta (4-8Hz), alpha (8-13Hz), beta (13-30Hz), gamma (30-70Hz), and high gamma (70-140Hz). Power spectra were calculated using the Welch method. Baseline LFP power was calculated by taking intervals of 1 second duration prior to each trial block, beginning 2 seconds before the start of the block and averaging all these pre-trial baselines. LFP power was baseline corrected by dividing the signal by this averaged baseline.

Electrophysiological features were calculated based on signals from individual contacts. Power within each frequency band was calculated for each contact. Network-level values were obtained by averaging features across electrodes assigned to the same functional network within each participant. Subject-level estimates were then averaged across participants for visualization purposes; SEM was calculated across patients. For decoding analyses (described below), each channel (and frequency band) was a separate feature.

For our analysis evaluating network states for high vs low creativity, we split the behavior data into thirds, and compared the LFP activity in different frequencies for the highest-scored third vs the lowest-scored third of each patient’s responses.

When comparing LFP power to baseline (or comparing AUT vs. BAT power), we computed patient-level difference waveforms by calculating the difference between power (across trial types or relative to baseline) at each time bin. Cluster-based permutation testing was performed across patients using a one-sample sign-flip procedure. At each time bin, the power difference was tested against zero using a one-sample *t* statistic. Adjacent time bins exceeding a threshold of *p* < 0.05 were grouped into clusters, with cluster mass defined as the summed absolute *t* statistic within each cluster. Statistical significance was assessed using a Monte Carlo null distribution generated by randomly flipping the sign of each patient’s power difference across 10,000 permutations. For each permutation, we noted the maximum cluster mass across the tested window. Observed clusters were assigned corrected *p*-values based on the proportion of permutations in which the maximum null cluster mass exceeded the observed cluster mass.

For completeness, we also report the AUT-BAT and AUT-OCT differences in power pooled across all patients and across the entire trial duration (all time bins in both the stimulus and response windows). However, we also directly compared AUT and BAT power for each participant, averaged across stimulus window time bins (two-sided paired *t*-tests). Meanwhile, we used a linear mixed effects model (LMM) from the Python statsmodels library^65^ to compare AUT vs. OCT LFP power in different trial windows—early stimulus (0-5 seconds), late stimulus (5-10 seconds), and response (10-15 seconds).

To quantify inter- and intra-network synchrony, we quantified the phase locking value (PLV) over time between contact pairs (using a 2 second sliding window), based on the instantaneous phase of LFP power at each recording site. PLV in each window (*w)* was calculated as: 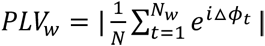, where △ *φ_t_* is the circular difference between instantaneous phase at time *t* across a pair of contacts, N is the number of timepoints in the window. To examine the relationship between phase locking and trial-level originality, we used a mixed effects model to account for non-independent data within-patient. We also used OLS to model the relationship between coherence and originality at the patient level, to inspect individual differences.

### Decoding

This paper interprets successful decoding in a given network as reflecting that this network contains information relevant to cognitive state or creative output.

K-fold cross validation was performed within-subject for linear decoders and for supervised CEBRA models. For all decoding analyses, except for decoding based on the self-supervised CEBRA embeddings, we held out sets of trials for cross-validation. In other words, both linear classifiers and supervised CEBRA models were trained on a subset of trials for each subject. (For 4-fold cross validation, each dataset was divided into 4 test-train splits based on trial index; timepoints within the same trial were kept together.) Linear classifiers were then tested on the held out trials, while trained CEBRA models were used to transform the held-out data into embeddings.

#### Linear classification

We used Python scikit-learn software^66^ to train a support vector machine (SVM) with a linear kernel to classify trials as belonging to AUT or BAT types, based on LFP power at different frequency bands across the time course of the trial, or based on firing rates of single cells. To control for discrepancies in electrode coverage across subjects and networks (especially between DMN and DAN—the two networks we most often directly compare), we ran the following three analyses:

1. Checked for the presence of a correlation between number of contacts in each network and decoding performance.
2. Reran decoding from DMN only in patients that also had coverage in DAN.
3. Reran decoding for DMN and DAN, subsampling contacts so that we were decoding from the same number of contacts for each patient. Contacts were subsampled randomly in 10 iterations, and results were averaged across the iterations prior to statistical comparison of DMN and DAN to avoid artificially inflating significance through higher number of iterations.

#### Nonlinear dimensionality reduction and decoding

We used CEBRA to identify low-dimensional latent embeddings from the high dimensional data^35^. For LFP data, we generated 5-dimensional embeddings, meaning that information from different channels and frequency bands was reduced into 5 dimensions. We confirmed that this value (5 dimensions) was a reasonable choice by generating creativity-related embeddings using all numbers of latent dimensions from 0-10 and inspecting creativity decoding performance (cross-validated) based on these embeddings (Supp. Fig. 8b). Single unit data had lower dimensionality already, so we generated 3-dimensional embeddings instead.

We fit independent CEBRA models on the data for each patient, then used the trained model to transform the data into the low dimensional embeddings. We then aligned the embeddings across all patients with Procrustes alignment. We used 4-fold cross validation to train the supervised CEBRA models: we randomly split the data into 4 folds, preserving temporal structure (timepoints within the same trial were not separated; only whole trials were held out). Train/test splits for each subject were chosen prior to training the models. We then trained the CEBRA models for each patient, holding out each fold (test set) while training on the remaining data (training set). For each fold, we then used a K-nearest neighbors (KNN) classifier to predict trial labels from the embeddings (concatenated across patients for each fold). For self-supervised analysis, no data was held out during CEBRA training, since these models never had access to labels, but we held out one subject at a time for the KNN decoding based on embeddings (where the 5 embedding dimensions constituted the features).

Since we aligned embeddings across patients and only ran the decoder once per validation fold, we first simply used a 1-sample t-test to compare the performance on all cross-validation folds to chance level. Traditional permutation tests with CEBRA models were not feasible, given the size of the data and the required computing power. Instead, to confirm that our model validation was correct (i.e., no “data leakage” occurred during decoding) and that the model was in fact learning real patterns in the data, we also trained a CEBRA model on data with shuffled labels for each validation fold and confirmed that these models could not fit the shuffled datasets and decoding failed.

When decoding creativity score, we discretized and normalized the continuous score into 10 bins, so we defined chance level in this situation as 10%. For visualization (e.g., Fig. 4b) and for the linear regression to define a coding axis, we trained the CEBRA models on the continuous score, normalized from 0 to 1 to improve consistency across patients.

For the analysis that clustered network activity, we used network identity as a supervisory signal for a multi-subject CEBRA model, and only trained the model on AUT trials. For network clustering, all contacts were kept separate, and dimensionality reduction was performed across trials.

### Identifying creativity coding axis

A CEBRA model was trained on all patients’ datasets, with continuous normalized creativity score as a supervisory signal, resulting in 5 dimensional embeddings for LFP data and 3 dimensional embeddings for single unit data. Then, we fit a linear regression model and used the model weights to compute the creativity coding axis as a 5- or 3-dimensional unit vector that best explains the variance in the embedding. We then projected the embedding onto this axis (dot product of embedding and coding axis vector), yielding a 1-dimensional representation of creativity values along the coding axis. The same process was used to project data onto the trial type coding axis.

Note that using normalized continuous creativity values as labels (i.e., eliminating individual differences in behavior between patients) improves the model fit (DMN: *R^2^*=0.86; DAN: *R^2^*=0.73) and correlations (DMN *r*=0.93; DAN *r*=0.86) relative to raw originality scores (DMN: *R^2^*=0.77, *r*=0.87; DAN: *R^2^*=0.66, *r*=0.79), but the same pattern remained, with the most descriptive coding axis being within the DMN.

### Single unit analysis

Spike sorting was performed using Offline Sorter (Plexon, Dallas, TX). Initially, the signal threshold was set to −3.5 RMS. Next, an automatic spike detection was applied with a T-Distribution Expectation Maximization sorting method (parameters: degrees of freedom = 4, # of expected units = 3). Then, the classification of waveforms was followed by manual curation with each microelectrode channel being visually inspected for additional units that went undetected during the automatic sorting step.

Units came from the following regions: 47 in medial temporal lobe, 34 in ventromedial prefrontal cortex, 13 in dorsomedial prefrontal cortex, 20 in middle cingulate cortex, and 8 in posterior cingulate cortex.

To determine trial type selectivity for the 3 trial types (AUT, BAT, and OCT) we performed one-way ANOVA (FDR-corrected for multiple comparisons) to identify cells with significant selectivity. We then performed paired t-tests to find which cells has a specific preference to a single trial type compared to both others: for example, a cell exhibiting responses characterized by AUT > OCT and AUT > BAT would be considered an AUT enhanced cell. This analysis was done for the 6 patients who completed all three trial types. For the 10 patients who only completed AUT and BAT, we compared AUT and BAT responses directly with paired t-tests. For CEBRA decoding of AUT vs. OCT trial types, we used 6-fold cross validation to train the CEBRA models, due to the smaller dataset.

### Statistics

Nonparametric statistical tests were used when comparing across networks due to the variable coverage. We report *t* statistics with degrees of freedom in parentheses. False detection rate (FDR) correction for multiple comparisons was applied for tests considering the 7 cortical networks, as these tests always had 7 comparisons. For these comparisons, all reported *p*-values (but not *t*-values) are corrected for multiple comparisons. FDR was also applied when determining selectivity of single unit responses, since we had to do a comparison for each of the 127 cells in the AUT-BAT contrast (or 85 cells for the AUT-BAT-OCT contrast). All t-tests and nonparametric null hypothesis tests were two-tailed unless otherwise noted. All reported correlation coefficients *r* come from Pearson correlation tests. The goodness of fit of the linear regression that identified the coding axes was quantified as the coefficient of determination (*R^2^*).

When directly comparing correlations or goodness of fit of specific networks (e.g., DMN vs DAN), we also re-did the comparison including only subjects that had coverage in both networks, and saw no difference in the results. Similarly, we reran the cognitive state decoding for DMN in only those patients than also had coverage in DAN and found equivalent decoding performance: for this subset of DMN datasets, AUC = 0.78 (whereas for the full set of DMN datasets, as reported in Supp. Fig. 2c, AUC = 0.77).

## Data availability

Processed behavioral and neural data underlying the figures and supplementary figures are available in Zenodo at 10.5281/zenodo.24058884. Raw behavioral and electrophysiological data are not publicly available because they contain sensitive or potentially identifying information and are subject to participant-consent and institutional restrictions. Deidentified data may be shared with qualified researchers for non-commercial research following review by the corresponding author and execution of a data-use agreement. Data requests should be directed to the corresponding author (Ben Shofty) and will be reviewed within 2 weeks.

## Code availability

Custom analysis code is available at https://github.com/ShoftyLabUtah/CreativityCode and is archived in Zenodo at 10.5281/zenodo.24058884.

## Acknowledgements

We thank the labs of Elliot Smith, Ben Shofty, Cory Inman, and Shervin Rahimpour for feedback and discussion; Alex Price for help with electrophysiology recordings; Justin Campbell for LFP preprocessing instruction; Veronica Zarr for help with patient consent; Jack Bowler for analysis advice and comments on the manuscript; and the patients for participating in experiments.

## Author contributions

B.S. and RR conceptualized the project. B.S. and E.H.S supervised the project. B.S., R.R., E.B., E.H.S., R.E.B., and S.R. designed the experiments and critically reviewed the methods. B.S., S.A.S., R.R., E.H.S., R.E.B., and T.S.D. developed the overall research strategy. B.S. and S.R. performed the surgeries. R.R., R.L.C., M.R.L., and T.S.D. collected and analyzed electrophysiological and behavioral data. R.R., E.H.S., and R.L.C. contributed to data preprocessing pipelines and software development. B.S. and R.E.B. provided conceptual input on cognitive paradigms and creativity metrics. R.R., and R.L.C performed the single-unit analyses. R.R., E.H.S, and B.S. wrote the original draft of the manuscript. E.H.S., B.S., R.L.C., T.S.D., S.R., E.B., R.E.B., and S.A.S. provided feedback for editing the manuscript. E.H.S. provided critical guidance on electrophysiology and clinical implementation. All authors contributed to data interpretation and provided critical revisions to the manuscript.

