## Supplementary Materials for "Originality in creative ideation evolves along nonlinear axes in the human brain"

**Supporting Information**

**Supplementary Results**

To further address the concern that weaker DAN decoding might reflect sparse DAN coverage, we added a contact-level sensitivity analysis. For each contact, we decoded trial type using frequency-by-time as features, and computed the cross-validated decoding AUC. We then modeled contact-level AUC as a function of DAN (vs. DMN) membership while accounting for participant-level clustering. DAN contacts showed lower decoding performance for the AUT-BAT contrast than DMN contacts in an ordinary least squares model with participant-clustered standard errors ($\beta$=-0.074, 95% CI [-0.136, -0.013], $p$=0.018). Further, to ensure this result was not driven by a single participant, we performed a leave-one-subject-out robustness analysis, repeating the decoding while holding out DAN data from one subject at a time. The DAN coefficient remained negative in all leave-one-subject-out iterations (ranging from $\beta$=-0.051 to $\beta$=-0.091). Thus, the weaker DAN decoding cannot simply be attributed to fewer DAN contacts, nor to DAN contacts in any single participant. We have added these results to the revised manuscript. The AUT-OCT decoding revealed a similar result ($\beta$=-0.051, 95% CI [-0.098, -0.0036, $p$=0.035). Although these analyses cannot entirely rule out the possibility that denser/different DAN coverage could reveal stronger DAN-based decoding performance, they do suggest that our results are not simply an effect the number of DAN contacts. We used ordinary least squares (OLS) regression (from the statsmodels library) for the contact-level sensitivity analysis (comparing DMN and DAN AUCs for contact-level decoding of trial type). For both the contact-level decoding and population linear decoding, we used SVM for the AUT-BAT contrast and linear discriminant analysis (LDA) for the AUT-OCT contrast.

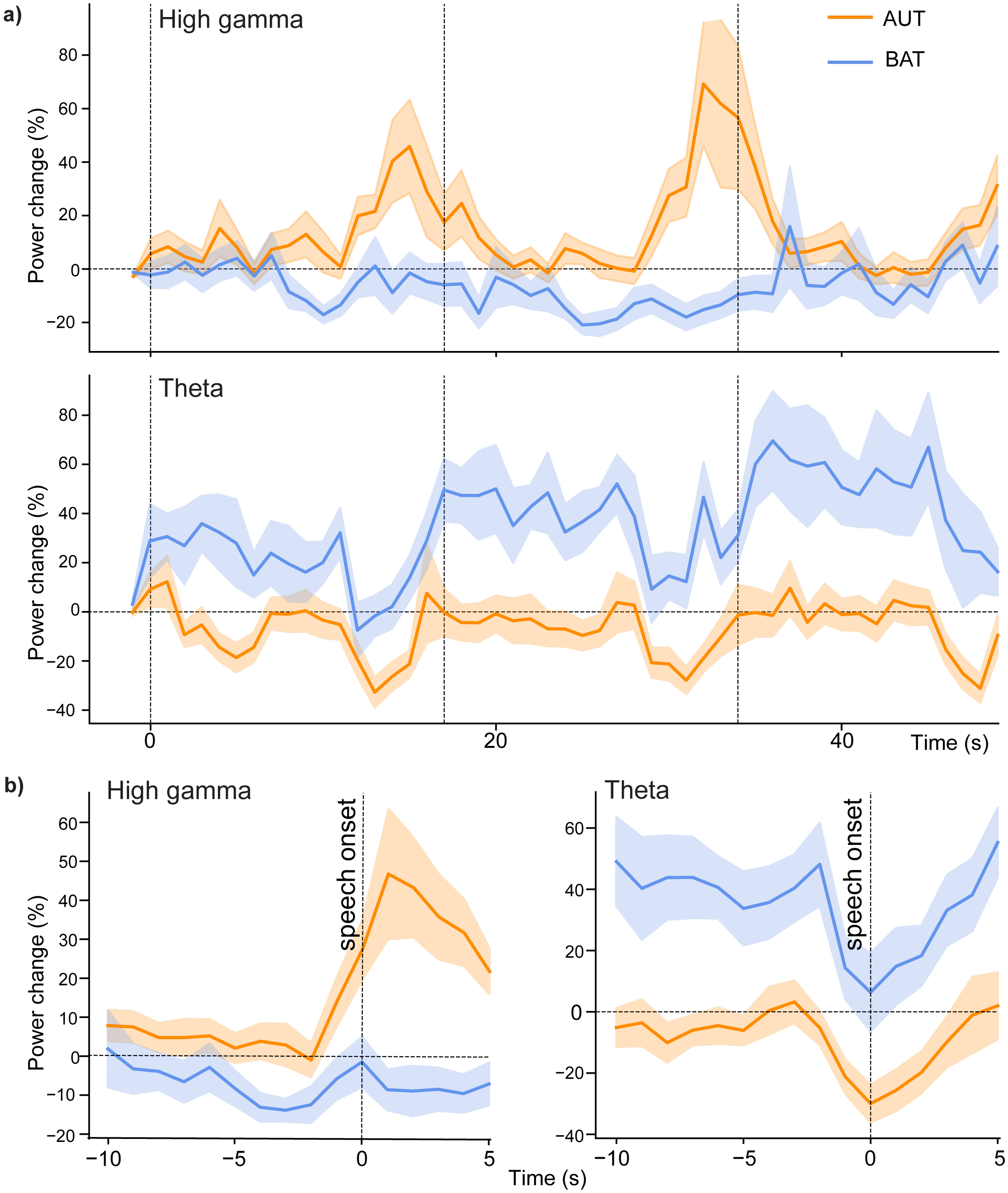


**Supplementary Figure 1. a)** Time course of high frequency (high gamma; 70-140 Hz) and low frequency (theta; 4-8 Hz) power for the alternate uses task (AUT; orange) and basic arithmetic task (BAT, blue), for the entire block of three trials relative to a pre-block baseline. Each vertical dashed line is the start of a new trial (stimulus onset). Power change is calculated relative to a baseline computed as the average of all intervals 2 – 1 seconds before the start of each block. **b)** Time courses of LFP power (high gamma, left; theta, right) aligned to speech onset, rather than to trial events. Shading indicates standard error. n=15 subjects.

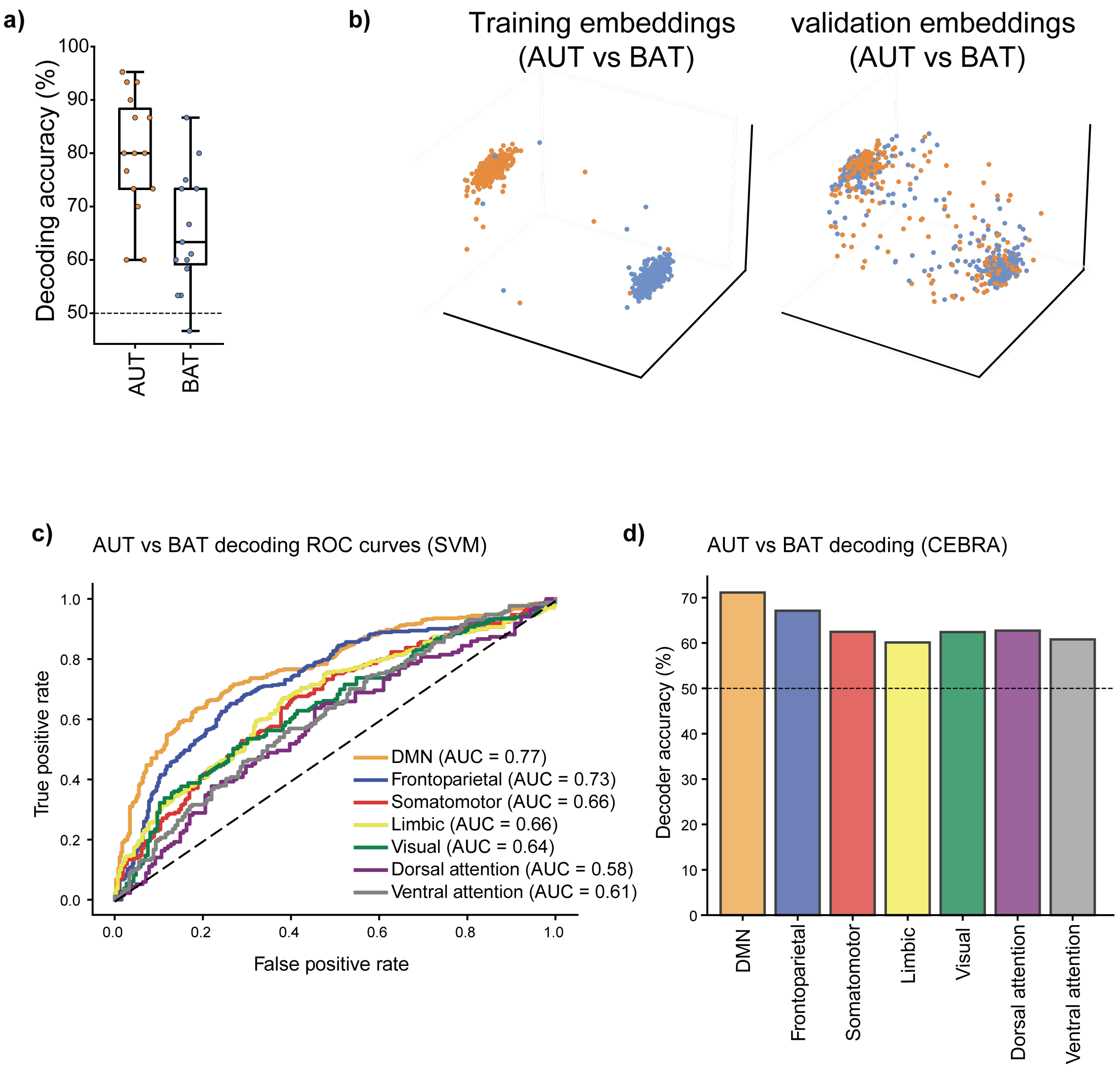


**Supplementary Figure 2.**
**a)** Linear SVM cognitive state decoding accuracy for AUT ($t$=10.3; $p$=6.3 × $10^{-8}$; two-sided 1-sample $t$-test) vs. BAT ($t$=5.7; $p$=5.9 × $10^{-5}$) trials. Each dot is the decoding performance for a single subject. n=15 subjects.
**b)** Visualization of trial-type separation, based on low-dimensional embeddings from the training and testing set of 1 cross-validation fold. Left: embeddings generated by CEBRA models trained on ¾ of the trials for each patient to separate AUT (orange) and BAT (blue) trials. Right: Held-out data transformed into the same latent space.
**c)** ROC curves to quantify trial type decoding performance of a linear SVM, across networks. Corresponds to result shown in Main Fig. 3d. n=15 subjects.
**d)** Trial type decoding performance based on each network's latent embeddings, as generated by CEBRA models trained to separate AUT and BAT trials.

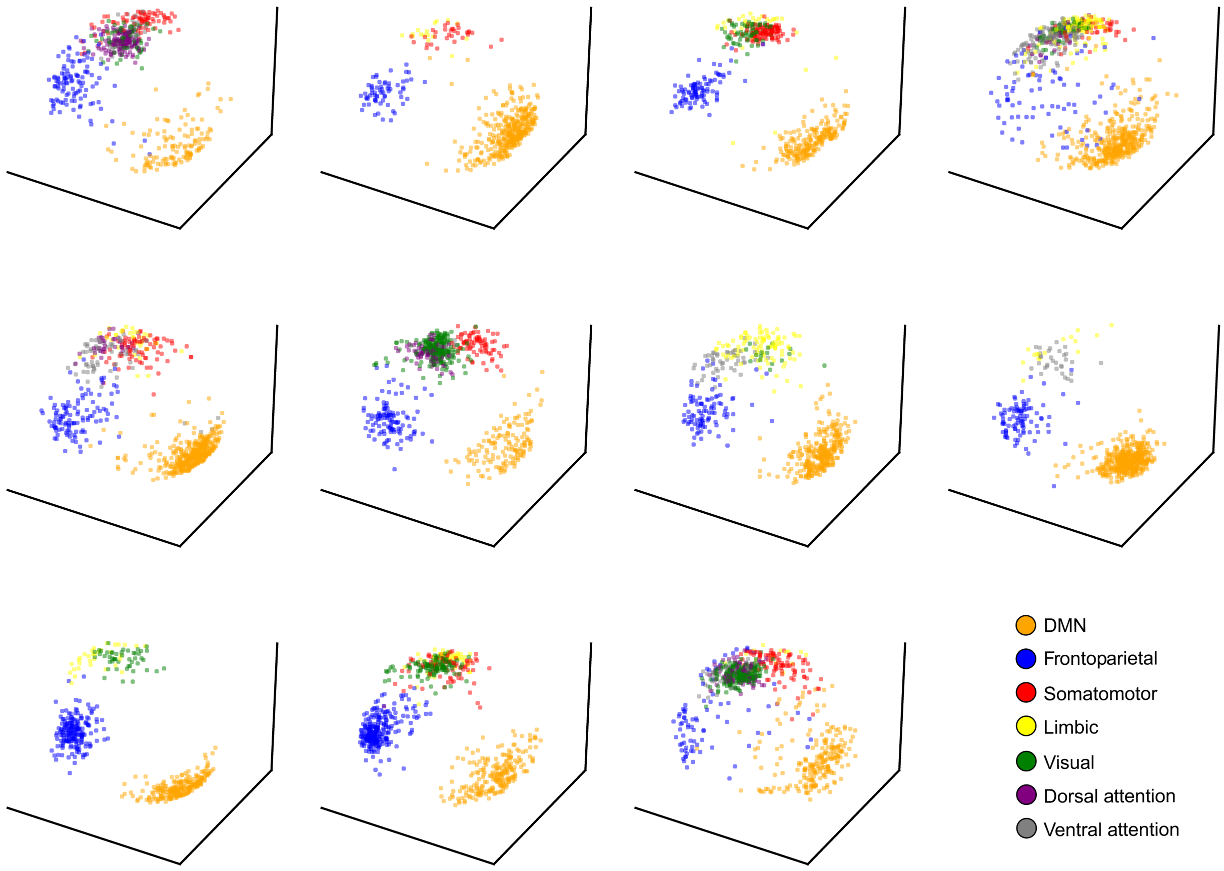


**Supplementary Figure 3.** Clustered network activity for individual subjects. Dimensionality reduction using CEBRA trained to separate activity of different networks, as in Main Fig. 3e, but for individual patients (first 11 patients shown).

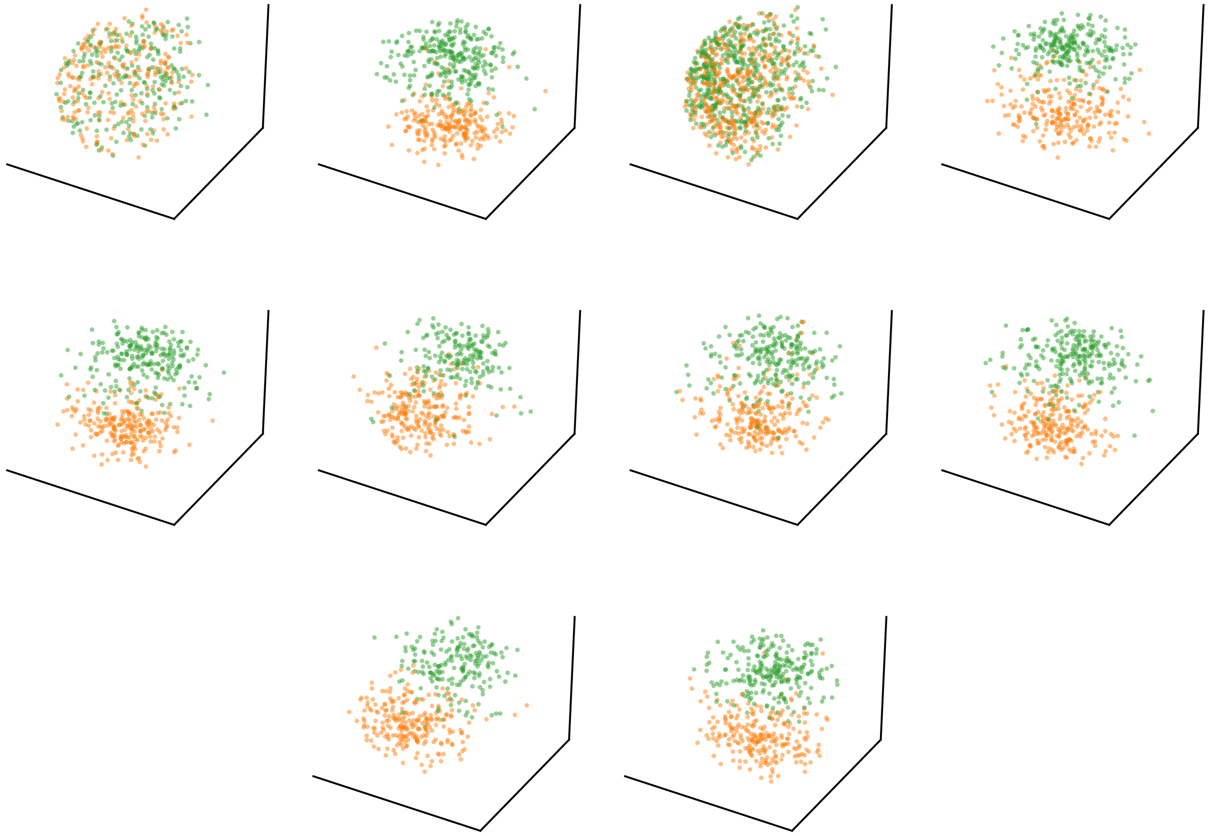

**Supplementary Figure 4.** CEBRA embeddings generated by models trained with AUT and OCT labels as the supervisory signal, illustrating qualitative separation between these two trial types in the DMN. Each panel is the embedding for a single patient (n=10 subjects). Decoding on embeddings generated from held out data yielded 59.18% decoding accuracy ($p$ = 0.0085; two-tailed one-sample $t$-test; FDR corrected for multiple comparisons across 7 networks).

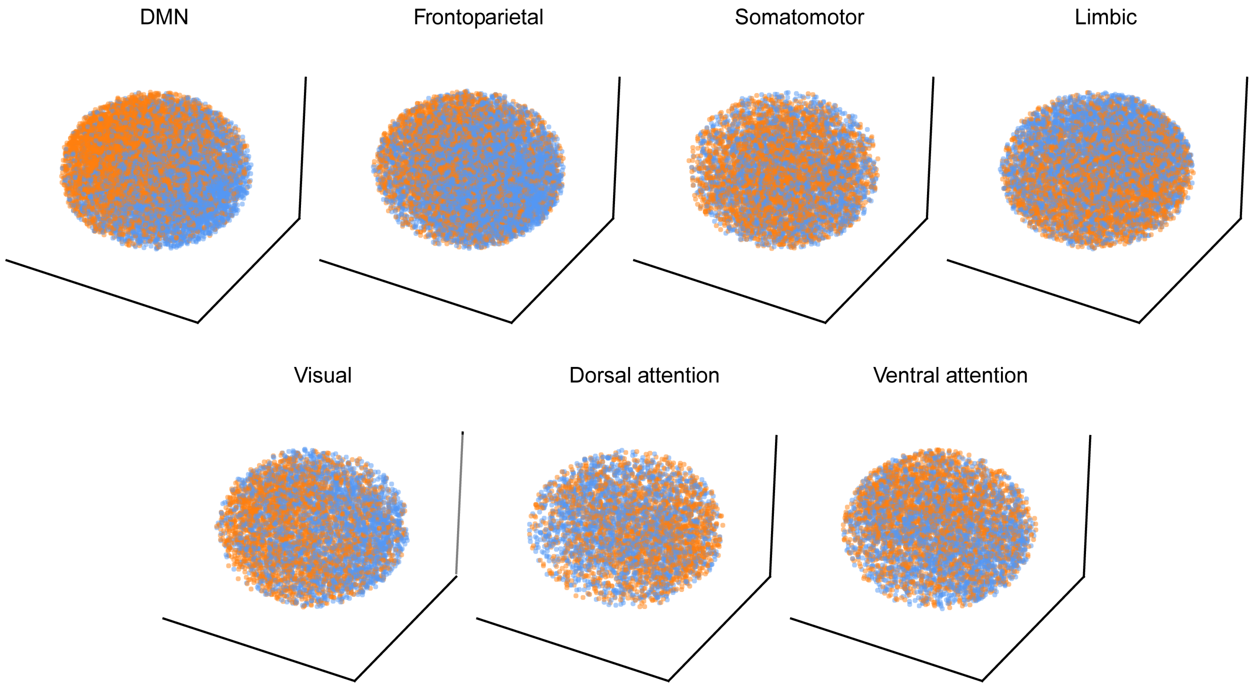


**Supplementary Figure 5.** Default mode network activity patterns intrinsically represent cognitive state. Latent embeddings generated by self-supervised CEBRA models (only time-contrastive training; no behavior labels to supervise training) for each network. Each dot is 1 timepoint. Embeddings are color-coded by trial type. AUT trials are orange and BAT trials are blue.

.

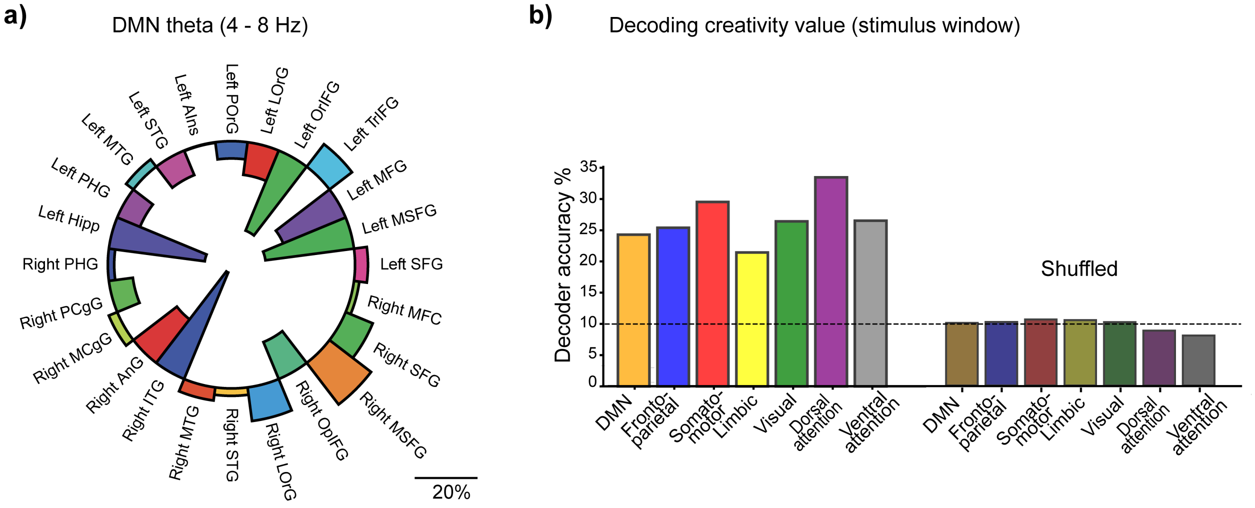


**Supplementary Figure 6.**
**a)** As in Main Fig. 4a, but for theta LFP power, comparing most creative – least creative trials.
**b)** Decoding creativity level based on the latent embeddings, as in Main Fig. 4c, but for only the stimulus window. Bars on the right show decoding based on models trained with shuffled labels.

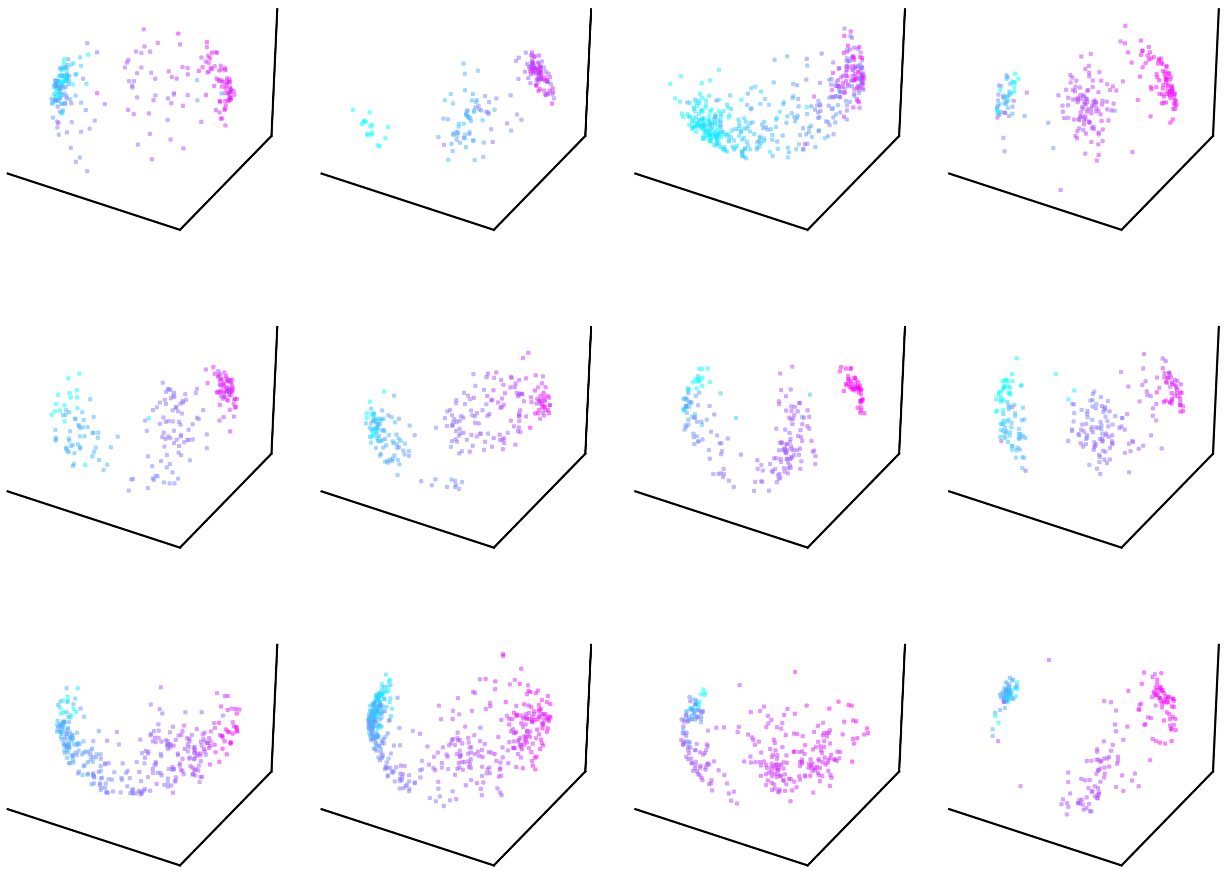

**Supplementary Figure 7.** Latent embeddings for originality level based on a CEBRA model trained on DMN data from all subjects; embeddings shown for individual subjects (first 12 patients).

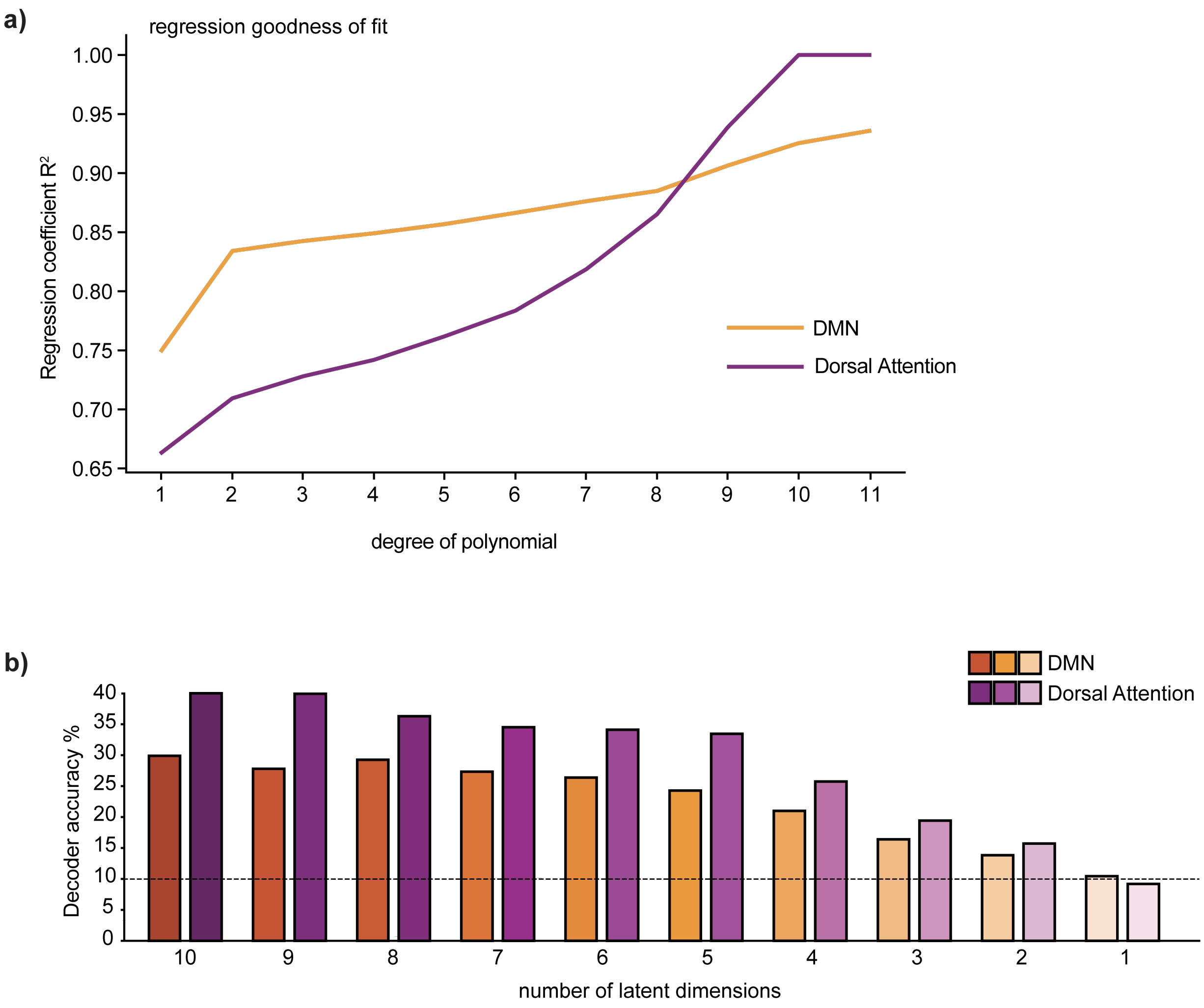


**Supplementary Figure 8.**
**a)** Goodness of fit of polynomial regression for DMN (orange) and DAN (purple), across different degrees of the polynomial.
**b)** Decoding performance (creativity level) based on DMN and dorsal attention network embedding, for different numbers of latent dimensions. Performance increases with increased number of dimensions but starts to plateau after 5 dimensions. When reducing to a single dimension, decoding performance falls to chance for both networks. Dorsal attention network yields higher decoding performance for most numbers of dimensions.

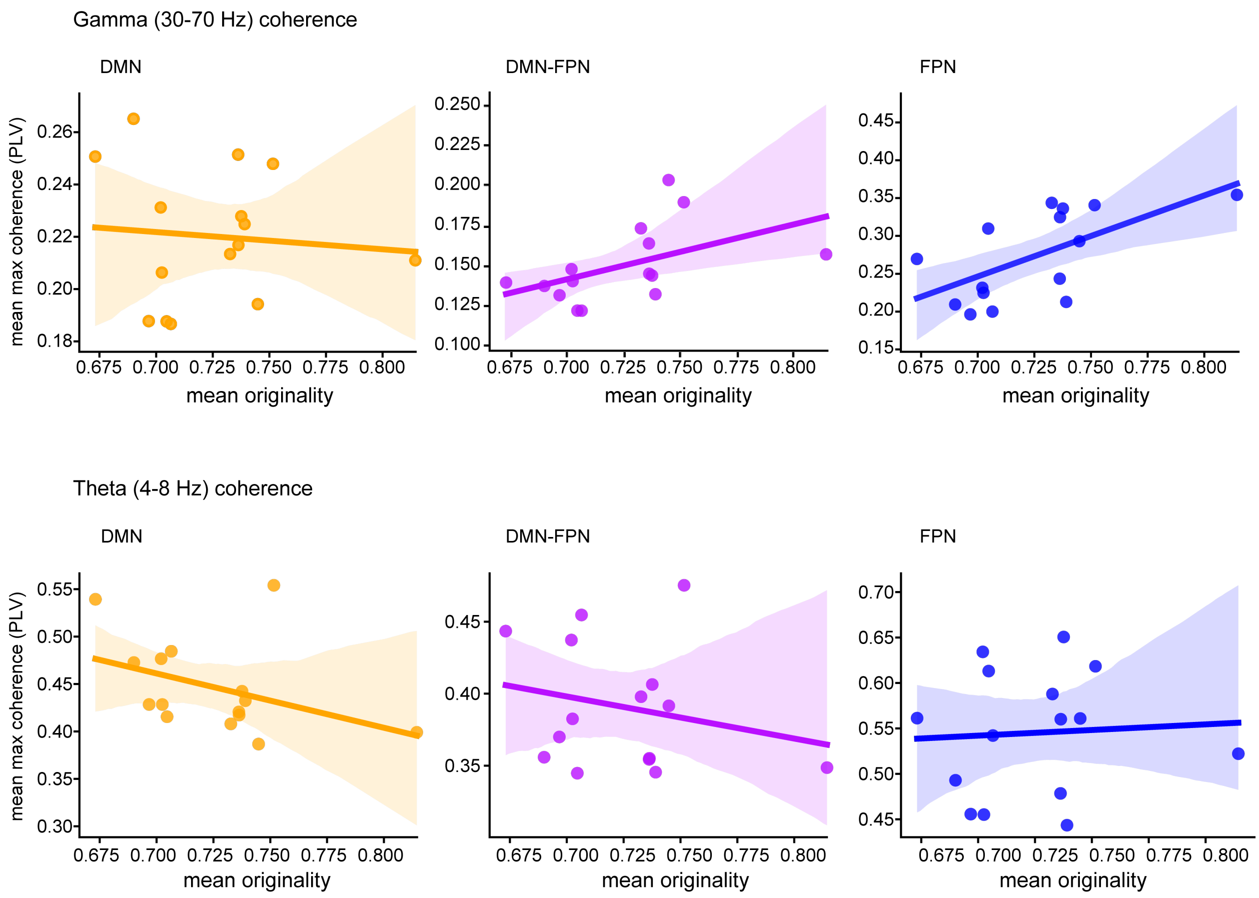


**Supplementary Figure 9**. Patient-level relationship between originality scores and peak phase coherence within and between DMN and FPN for gamma (top; 30-70 Hz) and theta (bottom; 4-8 Hz). Data from the stimulus period of the task (n=15 subjects). Shaded area indicates 95% confidence interval.
